# Direct and indirect effects of spliceosome disruption compromise gene regulation by Nonsense-Mediated mRNA Decay

**DOI:** 10.1101/2024.12.27.630533

**Authors:** Caleb M. Embree, Andreas Stephanou, Guramrit Singh

## Abstract

Pre-mRNA splicing, carried out in the nucleus by a large ribonucleoprotein machine known as the spliceosome, is functionally and physically coupled to the mRNA surveillance pathway in the cytoplasm called nonsense mediated mRNA decay (NMD). The NMD pathway monitors for premature translation termination signals, which can result from alternative splicing, by relying on the exon junction complex (EJC) deposited on exon-exon junctions by the spliceosome. Recently, multiple genetic screens in human cell lines have identified numerous spliceosome components as putative NMD factors. Using publicly available RNA-seq datasets from K562 and HepG2 cells depleted of 18 different spliceosome components, we find that natural NMD targeted mRNA isoforms are upregulated when members of the catalytic spliceosome are reduced. While some of this increase could be due to widespread pleiotropic effects of spliceosome dysfunction (e.g., reduced expression of NMD factors due to mis-splicing of their mRNAs), we identify that AQR, SF3B1, SF3B4 and CDC40 may have a more direct role in NMD. We also test the hypothesis that increased production of novel NMD substrates may overwhelm the pathway to find a direct correlation between the amount of novel NMD substrates detected and the degree of NMD inhibition observed. Finally, similar transcriptome alterations and NMD substrate upregulation are also observed in cells treated with spliceosome inhibitors and in cells derived from retinitis pigmentosa patients with mutations in *PRPF8* and *PRPF31*. Overall, our results show that regardless of the cause, spliceosome disruption upregulates a broad set of NMD targets, which could contribute to cellular dysfunction in spliceosomopathies.

**AUTHOR SUMMARY:** During gene expression, a complex cellular machine known as spliceosome removes extraneous non-coding sequences from precursor RNAs to produce messenger RNA (mRNA) with a contiguous code for protein sequence. To guard against splicing errors that may interrupt protein coding sequence, splicing is linked to a mRNA surveillance pathway known as nonsense mediated mRNA decay (NMD). In this work, we follow up on recent findings from multiple genetic screens that several spliceosome components are necessary for efficient NMD. Our analysis of transcriptomes of lymphoblast K562 cells depleted of 18 spliceosome factors show that NMD based regulation is compromised in cells lacking catalytic spliceosome proteins. Four of these spliceosome proteins may have a direct effect on NMD even though spliceosome disruption in general also causes other changes in gene expression that indirectly affect NMD. Our results suggest that defective NMD based regulation contributes to cellular dysfunction in spliceosomopathies, a collection of human genetic disorders caused by mutations in spliceosome factors.

## INTRODUCTION

Pre-mRNA splicing has a profound impact on mRNA substrates that are generated and translated into proteins. While alternative splicing generates multiple mRNA isoforms from a single gene to diversify the proteome, it also has the potential to impact open reading frame integrity and hence compromise protein expression. It is therefore not surprising that pre-mRNA splicing in the nucleus is functionally coupled to translation-linked nonsense-mediated mRNA decay (NMD) mechanism in the cytoplasm that identifies and rapidly degrades mRNAs containing premature translation termination codons (PTC) [1,2]. The influence of this connection has been widely documented in previous studies. For example, in *Saccharomyces cerevisiae* mutations in NMD components cause increased accumulation of erroneously spliced mRNAs [3] whereas, in human cell lines, numerous transcripts with disrupted open reading frames that are normally degraded and suppressed by NMD can be detected in the nucleus or among the pool of pre-translated mRNAs [4]. A particularly notable example of the functional connection between splicing and NMD is the process of alternative splicing coupled NMD (AS-NMD) where regulated alternative splicing of poison exons either introduce a PTC upon their inclusion or create one via frameshifting if the exon is excluded [1,2,5,6]. AS-NMD ties splicing and NMD together in a complex regulatory network used to fine tune gene expression, often via evolutionarily conserved poison exons in developmentally important genes [5,6].

Splicing and NMD are physically connected via deposition of the exon junction complex (EJC) 24 nucleotides (nt) upstream of exon-exon junctions during the catalytic steps of splicing [7]. During translation, presence of an EJC downstream of a stop codon is the most prominent sensor of premature termination (reviewed in [8–10]). Such 3’- untranslated region (UTR)-bound EJCs engage with the key NMD factors that include UPF1, UPF2 and UPF3, to mark the termination event as aberrant. Following sensing of aberrant termination, NMD is activated when UPF1 is phosphorylated by the SMG1 kinase to subsequently recruit SMG5, SMG6 and SMG7 proteins that either directly initiate mRNA degradation via SMG6-mediated endonucleolytic cleavage or by recruiting other mRNA decay enzymes through SMG5 and SMG7 [11–13].

Through their role in EJC deposition, many spliceosome components have been identified to play a direct role in NMD, thereby extending the connection between splicing and NMD. CWC22 and CWC27, two proteins recruited to the activated spliceosome, directly mediate recruitment and deposition of the EJC anchor, EIF4A3 [14–17], and thereby play an important role in NMD. AQR, also known as Intron Binding Protein 160 or IBP160, is another spliceosome protein implicated in EJC deposition through an as-yet unknown mechanism and has a documented role in NMD [18]. The structures of the spliceosome [19,20] that contain EJC (or pre-EJC) show other spliceosome proteins, such as EFTUD2, that come in direct contact with EJC subunits and may have a role in EJC assembly or deposition.

Two lines of evidence suggest that the connection between splicing and NMD may be more extensive than currently understood. First, human patients with mutations in several spliceosome components and the EJC proteins result in similar phenotypic effects. Mutations in pre-mRNA splicing components cause numerous disorders collectively referred to as spliceosomopathies [21], which can be classified into four broad categories: cranio-facial disorders, neurodevelopmental deficits, limb defects, myelodysplastic syndrome (MDS), and retinitis pigmentosa (RP), the latter of which is the most common.

Curiously, mutations in EJC subunits, notably the core protein EIF4A3, also cause cranio-facial disorders and neurodevelopmental defects [21–24]. Moreover, mutations in *EIF4A3* and *RBM8A*, another EJC core protein, like those in spliceosome components *SNRNPA* and *SF3B4*, lead to limb defects [24,25]. Second, several recent genetic screens for potential NMD factors have identified numerous spliceosome components among the top hits [26–29]. The identification of spliceosome proteins as potential NMD factors in these screens, which were performed using different NMD reporter RNAs and employed different candidate gene inactivation strategies, underscore the possibility that spliceosome components beyond CWC22, CWC27 and AQR may have yet to be determined roles in NMD.

To uncover the extent and modes of connection between the spliceosome and NMD, we analyzed publicly available RNA-seq datasets from human cell lines depleted of several spliceosome proteins, those treated with drugs that alter spliceosome function, and cells derived from patients with spliceosome mutation causing retinitis pigmentosa. In these samples, we quantified the changes in abundance of NMD-targeted transcripts and splicing patterns to identify changes to the transcriptome when spliceosome is disrupted. Our results show that depletion of many catalytic spliceosome components leads to an increased abundance of endogenous EJC-dependent NMD targets. In several cases, we also observe a similar increase in other non-canonical isoforms. These results indicate that depletion of spliceosome components broadly changes the transcriptome, resulting in upregulation of NMD-targeted transcripts through mis-splicing or reduction in NMD efficiency, or both. Interestingly, depletion of four spliceosomal components, AQR, CDC40, SF3B1 and SF3B4, like the knockdown of EJC core protein EIF4A3, causes higher upregulation of NMD targeted isoforms as compared to other non-canonical isoforms, suggesting that their effects on NMD could be more direct. All together, we show that disruption of pre-mRNA splicing has direct as well as pleotropic effects on gene expression that also results in increased expression of NMD targeted transcripts. Although the precise reason of this effect on NMD targets remains unresolved, altered levels of NMD regulated genes may contribute to the molecular phenotypes in spliceosomopathies.

## RESULTS

### Several components of catalytic spliceosome are overrepresented in genetic screens for novel NMD factors in human cell lines

In the search for novel NMD factors, a number of genome-wide genetic screens have been recently conducted in human cell lines using CRISPR-Cas9 mediated gene knockouts [26,28,29] or siRNA mediated knockdowns [27]. Even though these screens employed different NMD reporters and gene knockdown/knockout methodologies, a comparison of the top 200 factors identified in each of these four screens shows a substantial overlap among the factors that influence NMD (**Fig 1A**). A total of 691 genes are present in the top 200 list of the four screens. A functional protein association analysis between these 691 proteins using STRING [30] shows that gene ontology terms related to mRNA metabolism including pre-mRNA splicing and NMD are enriched in this set (**Table S2**). Further, spliceosome factors, as defined by the spliceosome database [31], constitute 170 of the 691 proteins (**Table S3**). Surprisingly, among a more stringent list of 65 proteins that are present in the top 200 hits of two or more screens, 43 are spliceosome factors that form an inter-connected network with known and novel NMD factors (**Fig 1B**).

**Fig 1.**
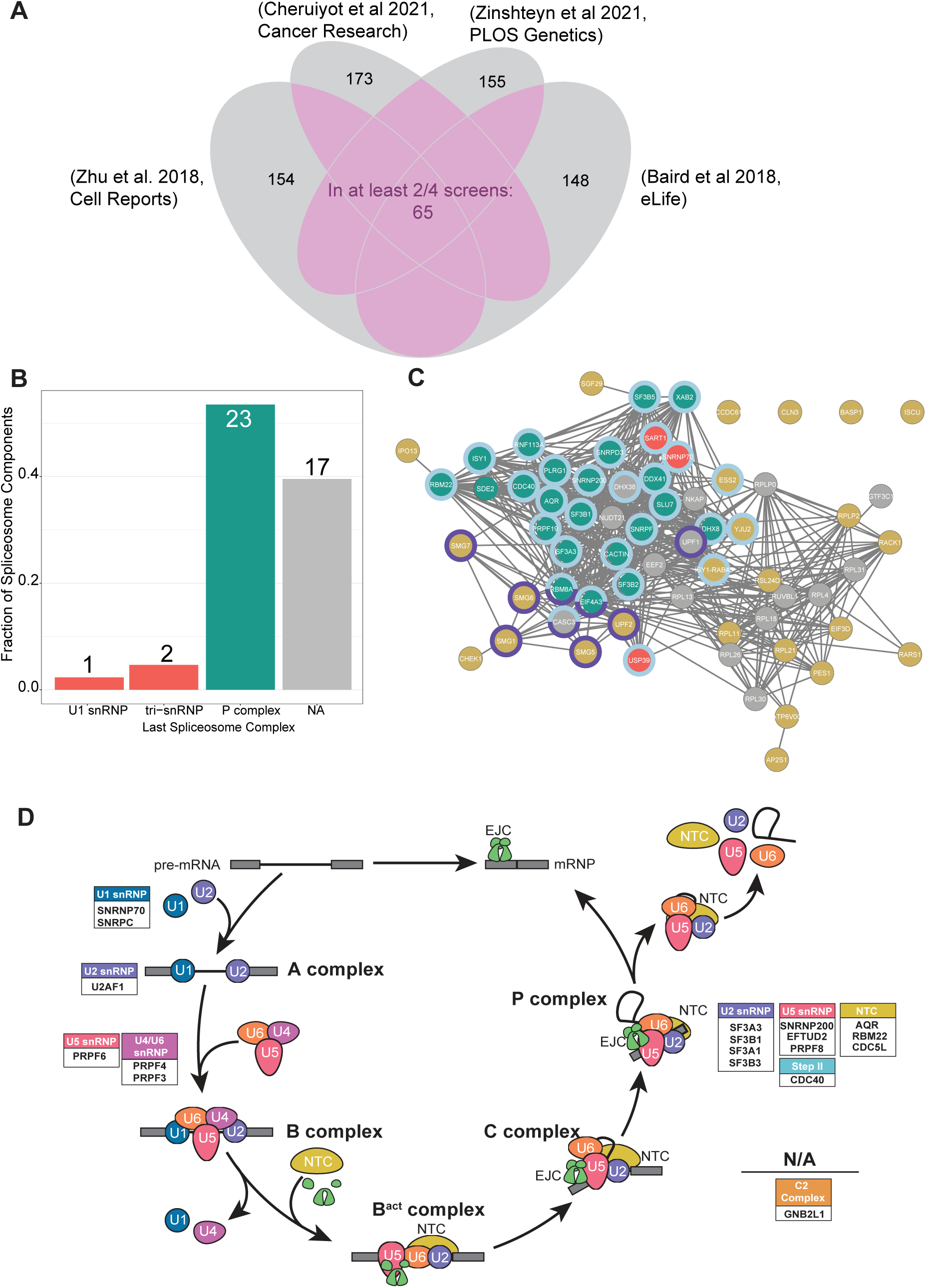
Spliceosome components identified in NMD factor screens are predominantly from catalytic spliceosome complexes. **A)** A Venn diagram showing overlaps between the top 200 hits in the indicated screens for NMD factors. 65 proteins were common in at least two of the four screens. **B)** A bar plot showing what fractions of spliceosome components identified in the NMD screens group into the spliceosomal complexes shown on the x-axis. The spliceosomal complex assignment was based on when a particular protein leaves the spliceosome, as defined by the spliceosome database. Early spliceosome components are colored red, catalytic spliceosome components are teal, and components with no annotated leaving time are grey. **C)** STRING network of protein-protein interactions of all factors identified in the top 200 of any two of the four screens (65 proteins from A). Nodes are colored according to when the protein leaves the spliceosome: red for early spliceosome components, teal for catalytic spliceosome components, grey for spliceosome components with no annotated leave point, and yellow for proteins that are not part of the spliceosome. Rings around nodes indicate gene ontology biological processes: RNA splicing (GO:0000398) in light blue; and nonsense-mediated decay (GO:0000184) in dark blue. **D)** The spliceosome components under investigation, grouped into splicing subcomplexes, are arranged around the splicing cycle where they leave the spliceosome.

The spliceosome undergoes a number of remodeling steps during the course of pre-mRNA splicing, meaning that not all components are present during all stages of the splicing cycle [32]. We categorized the identified spliceosome factors based on the last spliceosome complex they are associated with as defined in the spliceosome database (**Fig 1C and D**). Based on these classifications, we find that components of the P complex, the spliceosome that results immediately after exon-ligation step and before spliceosome disassembly, are overrepresented in NMD screens compared to their abundance in the spliceosome database (p-value < 2.2 x 10^-16^, Χ^2^ goodness of fit test) (**Fig 1B, C and D)**.

Given the tight coupling between pre-mRNA splicing and NMD via deposition of the NMD-enhancing EJC, some spliceosome components are expected to be enriched in screens for NMD factors. For example, CWC22, an integral spliceosome component with a direct role in recruitment and positioning of EIF4A3 on the 5’-exon within the catalytic spliceosome [14–16], is among or close to the top 200 hits in three of the screens (ranked 7, 233 and 379). Similarly, AQR, an NTC-related core spliceosome component with a documented role in EJC deposition albeit via unknown mechanism [18], is detected in the top 200 hits in three out of four screens. However, the identification of a surprisingly large number of catalytic spliceosome components in NMD screens suggests that connection between splicing and NMD is likely to extend far beyond the spliceosome components previously known to be involved in EJC deposition. Interestingly, no such enrichment is observed for components of early spliceosomal complexes that are involved in splice site recognition but leave before the catalytic steps (**Fig 1C**). Thus, the catalytic spliceosome appears to have a broader and yet to be fully appreciated impact on the proper functioning of NMD.

### Depletion of several catalytic spliceosome components increases abundance of endogenous NMD targets

To investigate the impact of spliceosome components on NMD, we decided to analyze data available from the ENCODE consortium as part of the ENCORE project [33–35] where individual proteins were knocked down or knocked out in K562 or HepG2 cell lines using siRNA or CRISPR-Cas9, respectively, followed by total RNA-Seq. Such RNA-Seq data is available from ENCODE for 16 of the 43 spliceosome proteins in the top 200 hits of at least 2 of the 4 screens. For two snRNPs, U1 and U5, only one component was present among these top hits. So, we included in our investigation additional members from these snRNPs (SNRNPC for U1; PRPF6, PRPF8 and EFTUD2 for U5) even though they were not among the 43 proteins shared among the NMD screens. For all these spliceosome component depletion datasets, we quantified gene expression at transcript level (using kallisto [36] and hg38 transcriptome build 109) and performed differential expression analysis (using DESeq2 [37]) between the depletion and wild-type (WT) datasets (**Table S4**). Any datasets where the depletion and WT replicates did not segregate in a principal component analysis or where the primary protein coding mRNA (as per the MANE select definition, [38]) of the protein being depleted was less than 2-fold downregulated were not analyzed further (**Fig S1**). Altogether we investigated 18 spliceosome components in total that met these criteria and represented among them are all snRNPs of the major spliceosome, as well as the nineteen complex (NTC) and NTC-related proteins (NTR) (**Fig 2A**). As controls, we used ENCODE RNA-Seq datasets from the same cell lines that were depleted for either key EJC proteins, MAGOH and EIF4A3, or the central NMD factor, UPF1.

**Fig 2.**
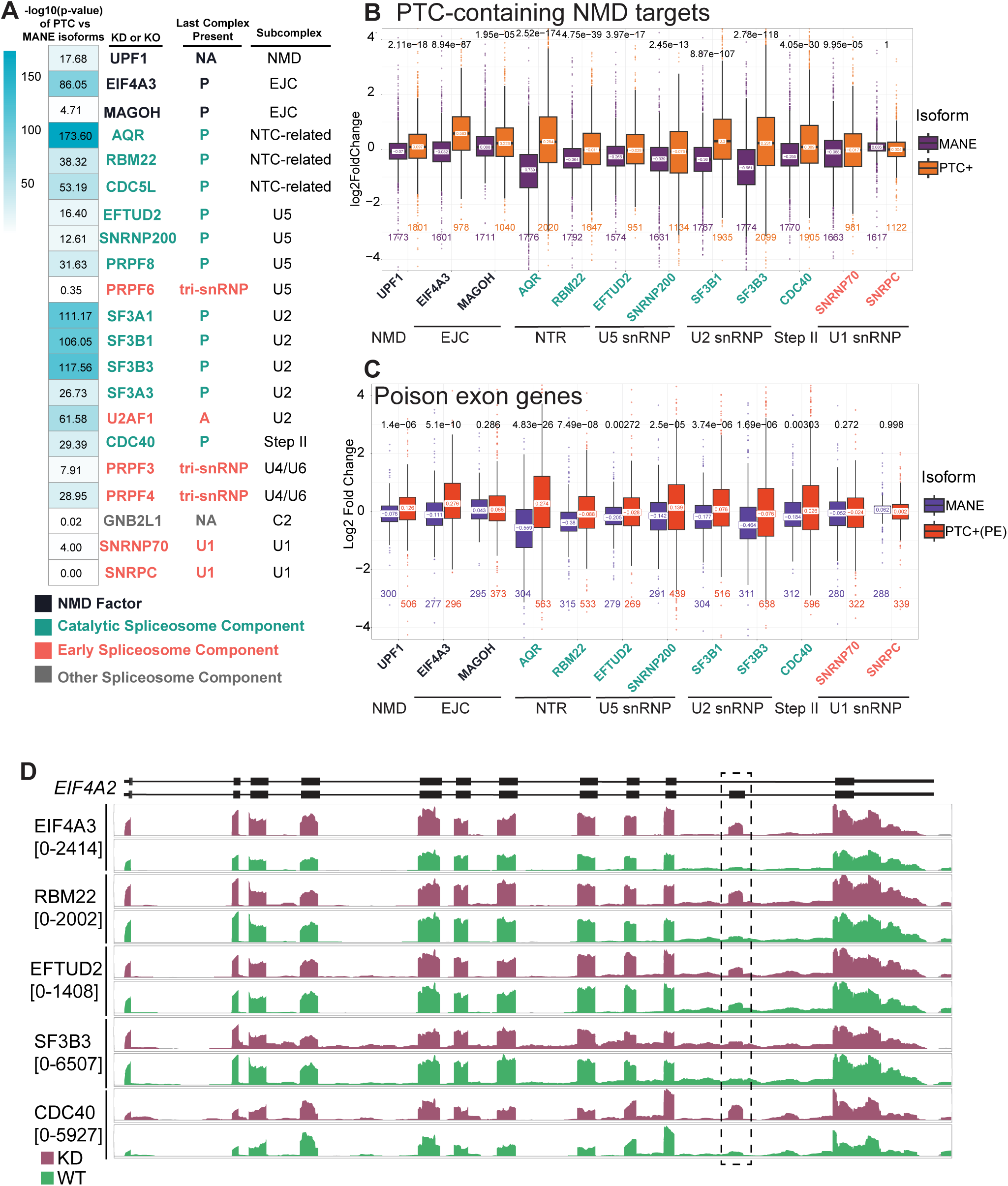
Depletion of many catalytic spliceosome components upregulates endogenous NMD targeted mRNAs in K562 cells. **A)** A heatmap on left displaying the-log10(p-value) of the Wilcoxon test comparing log2(fold change) of NMD-targeted as compared to MANE isoforms for all spliceosome component depletions tested. In the table to the right, spliceosome components are colored according to when they leave the spliceosome. Stable complexes the shown proteins are part of are also given. GNB2L1 and SNRNP70 knockdown datasets are from HepG2 cell lines while all others are from K562 cells. **B)** Boxplots displaying the log2(fold change) on the y-axis of MANE transcripts (purple) and NMD-targeted transcripts (orange) in the knockdowns indicated on x-axis. The number of transcripts in each group is indicated below the boxplot, the median of the boxplot is indicated on the boxplot, and the p-value of the Wilcoxon text comparing the two groups is above. Spliceosome component names are colored according to when they leave the spliceosome as in A. **C)** Boxplot showing the log2(fold change) of MANE transcripts (blue) and NMD-targeted transcripts (red) from genes with conserved poison exons. Boxplot and depletion annotations are as in B. **D)** Genome browser view showing distribution of reads mapping to *EIF4A2* in a representative wildtype (green) and knockdown (purple) replicates. The poison exon in *EIF4A2* along with mapping reads are indicated by the dashed box, and the scale of each pair of tracks is indicated below the knockdown name on the left. Shown at the bottom is the exon structure of MANE select and PTC+ transcripts (thin lines: introns, thick lines: exons).

To determine the effect of depletion of individual spliceosome components on NMD, we focused on endogenous transcripts that contain a termination codon at least 50 nucleotides upstream of an exon-exon junction, a set that we previously defined in Yi et al [39]. Such transcripts are targeted to NMD due to presence of an EJC downstream of a termination codon, which is thus regarded as premature termination codon (PTC) [39]. As not all PTC-containing transcripts undergo efficient NMD, we limited our analysis to only those PTC-containing transcripts (PTC+) that were previously shown to be upregulated upon combined depletion of SMG6 and SMG7 [12]. These PTC+ mRNAs are produced from ∼2,000 genes and therefore provide a broad measure of NMD activity. As control, we examined the effect of the depletions on the MANE select transcripts produced from the same genes. We find that, like the depletion of known NMD/EJC factors, reduced levels of all 11 catalytic spliceosome components tested lead to a higher median fold-change for PTC+ transcripts as compared to their corresponding MANE select isoforms (**Fig 2A, 2B** and **S2A**). Among these, the strongest effect on the PTC+ group is observed for AQR, an NTC-related protein previously reported to aid EJC assembly [18] Notably, depletion of two other NTC-related proteins, RBM22 and CDC5L, significantly increased PTC+ transcripts suggesting that NTC-related components beyond AQR may play a role in EJC-dependent NMD (**Fig 2A, 2B** and **S2A**). Interestingly, components of the SF3B (SF3B1, SF3B3) and SF3A (SF3A1) subcomplexes of the U2 snRNP also show a highly significant increase in median fold change for PTC+ transcripts as compared to the control group [(median fold-change for PTC+ versus MANE select transcripts – SF3B1: 0.3 vs - 0.36; SF3B3: 0.231 vs-0.661; SF3A1: 0.178 vs-0.355)] (**Fig 2A, 2B** and **S2A**). Notably, a mutation in SF3B1 that causes myelodysplastic syndrome was previously shown to upregulate endogenous NMD targets as well as an NMD reporter [28]. Our results suggest that the link between NMD target abundance and SF3 subcomplexes of U2 snRNP extends beyond SF3B1 (**Fig 2A, 2B** and **S2A**). Of the four U5 components tested, depletion of all but PRPF6 results in upregulation of the PTC+ transcripts compared to the MANE transcripts. Notably, PRPF6 is the only one of the U5 components tested that leaves before the spliceosome activation [31], further hinting that the effect on NMD as a result of spliceosome disruption could be tied to the two catalytic steps of splicing. Of the other U5 components, EFTUD2, which sits adjacent to EIF4A3 in the catalytic spliceosome and also engages with CWC22 [19], as well as SNRNP200 and PRPF8 shows a significant effect on the abundance of PTC+ transcripts (**Fig 2A, 2B** and **S2A**). Among the factors that leave before spliceosome activation depletion of only U2AF1, a U2 auxiliary factor and PRPF4, a U4 component, have a significant effect on PTC+ transcripts whereas both U1 snRNP components tested (SNRPC and SNRNP70), PRPF3, another U4 subunit, and GNB2L1, a C2 complex protein, have no or only a mild effect on NMD targeted transcripts (**Fig 2A, 2B** and **S2A**). We observed overall very similar trends in the effects of spliceosome component depletion when we limited our analysis to a specific class of PTC+ transcripts where splicing in (or inclusion) of PTC- containing poison-exons targets these mRNAs to NMD (**Fig 2C and S2B**). Notably, many poison exons are highly conserved and their inclusion is tightly linked to NMD-dependent transcript regulation [5], which can be critical for overall regulation of these genes and their functions [6]. A visual examination of read coverage of a well-documented specific poison exon, exon 11 of *EIF4A2*, shows that more reads map to the poison exons in the knockdown samples as compared to the WT cells (**Fig 2D**). Overall, we conclude that upon loss of many spliceosome components, particularly those of the catalytic spliceosome, endogenous NMD targeted transcripts accumulate at a higher abundance in K562 cells.

### Compromised splicing activity upon spliceosome component depletion leads to increased abundance of other non-canonical isoforms

Depletion of spliceosome components is expected to cause widespread changes in splicing, which in turn can impact multiple steps in gene expression. Hence, the RNA-Seq datasets from spliceosome component depletions will encompass changes in mRNA biogenesis (e.g., splicing), mRNA degradation (e.g., NMD), and all intermediate steps. To test if the increase in abundance of PTC+ transcripts is due to an effect on NMD or due to preferential generation of these isoforms via splicing, we first quantified global splicing changes in all depletion datasets using rMATS turbo v4.3.0 [40]. As expected, depletion of spliceosome components alters global splicing patterns, resulting in significant changes in annotated splice site usage and also producing novel splicing events (**Fig 3A**). Notably, in all knockdowns, the number of novel splicing events observed is greater than the number of annotated splicing events that change significantly (**Fig 3A**). Further, all the spliceosome depletions tested produce a similar number of novel splicing events, with the exception of AQR, which produces an even greater number of novel splicing changes. Among the annotated events, the depletion of the catalytic spliceosome members, as compared to the early spliceosome components tends to cause a bigger change in significantly altered annotated splice events (compare red circles to green circles in **Fig 3A**). The most common annotated splicing event that changes when a spliceosome component is depleted is exon skipping, accounting for roughly half of the altered annotated splice events in most samples tested (**Fig 3B**, top). Interestingly, among the novel splicing events we observe a dramatic increase in alternative 3’-and 5’-splice-site usage upon spliceosome component depletion (**Fig 3B**, compare yellow and green sections of the bars between annotated (top) and novel (bottom) events). This change could result from increased usage of weaker splice sites by the compromised spliceosome. When the combined effect of altered annotated and novel splicing events is considered on well-expressed genes (at least one transcript with TPM > 5 in wild-type cells), we find that a large fraction of genes (0.21 to 0.55) are subjected to alternative splicing upon spliceosome component knockdown (**Fig 3C**).

**Fig 3.**
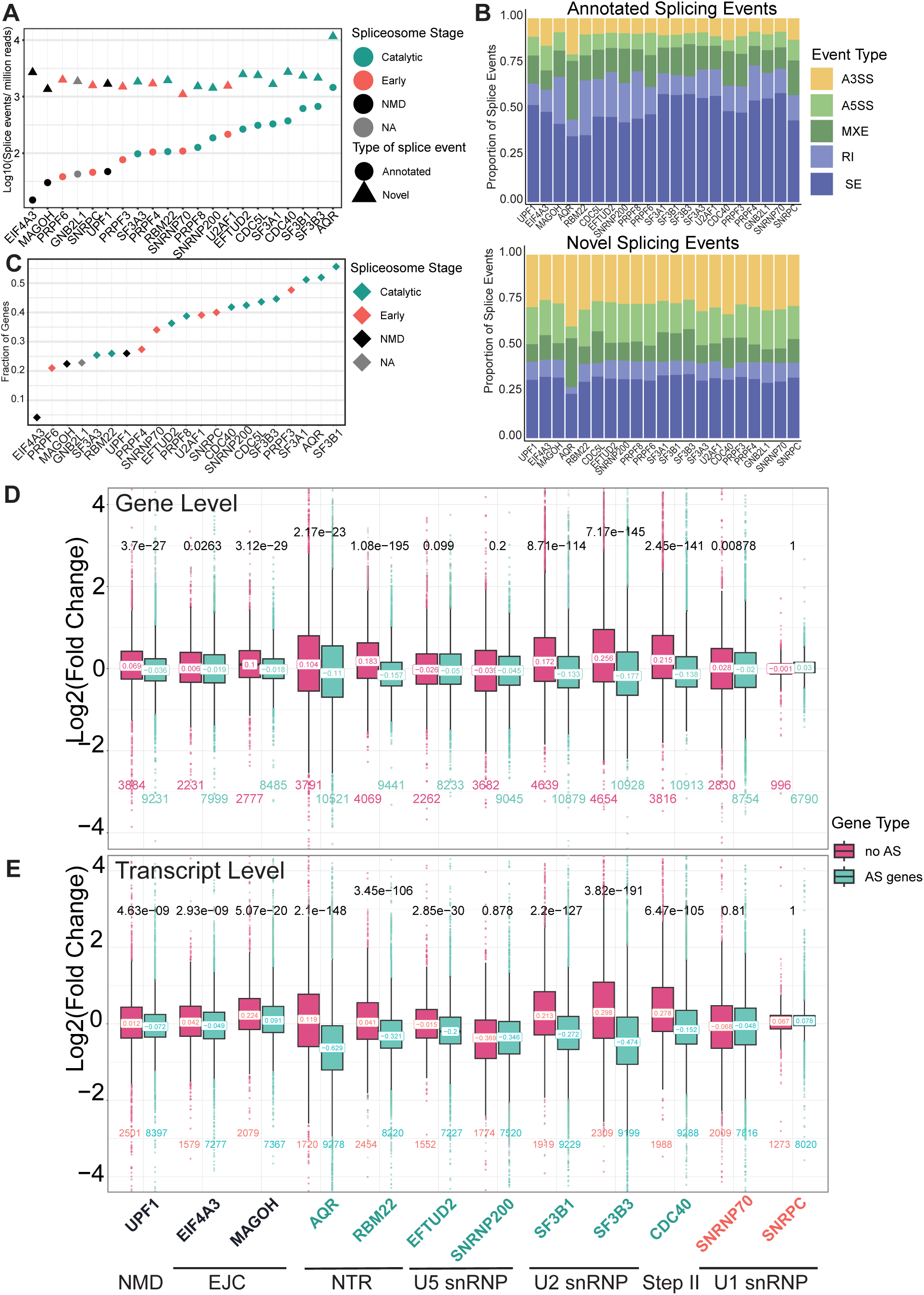
Widespread changes in annotated and novel splicing events upon spliceosome factor knockdowns reduce expression of the affected genes. **A)** A dot plot showing the count of splicing events (log10 transformed after normalizing to per million mapped reads) for annotated (circle points) and novel splicing events (triangle points). Points are colored according to when the components last leave the spliceosome. **B)** The proportion of significantly changing annotated (top) and novel splicing events (bottom) that are each splice type: alternate 3’ splice site (yellow), alternate 5’ splice site (light green), mutually exclusive exons (dark green), retained introns (light blue), and skipped exons (dark blue). **C)** The proportion of genes with length-scaled TPM > 5 in WT cells that have altered splicing patterns following the indicated KD. **D)** The log2(fold change) of genes that are (green) and are not (pink) undergoing altered splicing following spliceosome component knockdown. Comparisons are made at the gene level. **E)** The log2(fold change) of genes that are (green) and are not (pink) undergoing altered splicing following spliceosome component knockdown. Comparisons are made at the MANE transcript level.

It is expected that normal gene expression will be affected when spliceosome factor depletion causes widespread splicing alterations. Consistently, we observe that, upon depletion of a majority of catalytic spliceosome factors, genes that are subjected to alternative splicing (≥1 novel or significantly changing annotated AS event) show an overall downregulation as compared to genes with no detectable change in splicing (**Fig 3D** and **S3A**). In comparison, these effects are milder upon depletion of early spliceosome factors. We argued that the impact of altered splicing on gene expression under these conditions will be more apparent at transcript level as considerable changes in isoforms produced from a gene can still be masked in gene level estimates. Therefore, we performed transcript-level comparisons with a focus on canonical (MANE select) transcripts. It is conceivable that alternative splicing under such conditions could direct splicing of pre-mRNA away from the MANE isoform toward a different, potentially novel, isoform, thereby reducing the pool of MANE isoforms from that gene. Consistently, in almost all catalytic spliceosome factor depletion conditions, the MANE select isoforms from genes that show evidence of alternative splicing are reduced in their levels as compared to the MANE isoforms from genes where no significant or novel splicing changes are detected (**Fig 3E** and **S3B**). In contrast, effects on MANE isoform abundance of AS genes remain mild for early spliceosome component depletions. These results suggest that depletion of components of the catalytic spliceosome has a profound effect on mRNA isoform expression.

### Altered gene expression upon spliceosome component depletion also affects levels of non-NMD-targeted isoforms

The reduction in gene-level and MANE isoform expression upon catalytic spliceosome component depletion suggests that these conditions are also likely to affect the abundance of all isoforms produced from a gene, including the NMD targeted isoforms. Thus, we first compared the change in levels of the MANE select isoform versus other non-canonical isoforms for the genes that either are subjected to alternative splicing or not under the depletion conditions. We find that for genes with no evidence of alternative splicing the distributions of fold-change values for the MANE select and other non-canonical isoforms are comparable (i.e., not significantly different with a few exceptions among the catalytic spliceosome components) (**Fig S4A**). However, for the genes that undergo alternative splicing, the fold-change values of the non-canonical isoforms as compared to the canonical MANE select isoforms is significantly higher for all the catalytic spliceosome components tested (**Fig 4A** and **S4B**). Notably, knockdown of UPF1 or EJC factors and of the early spliceosome components show no or only a small difference in fold change distributions of MANE versus non-canonical isoforms from genes with or without alternative splicing. To separate the effects of alternative splicing and NMD on transcript levels, we attempted to identify genes that produce an NMD-targeted isoform but do not show evidence for alternative splicing upon spliceosome component depletion. However, we could find only a handful of such genes in most knockdowns, and thus could not perform any conclusive analysis.

**Fig 4.**
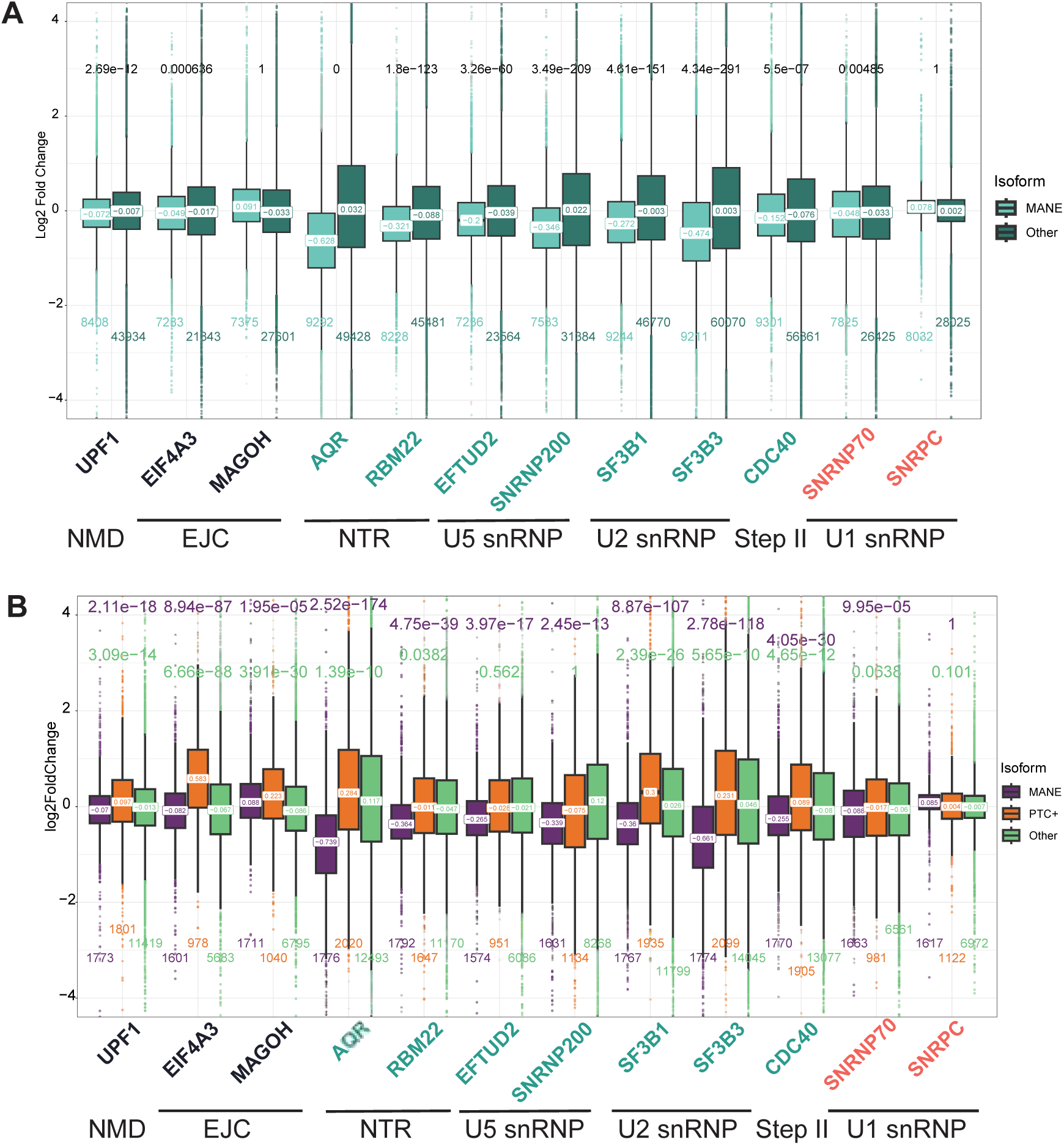
Transcript re-quantification after including novel isoforms reveals that all non-canonical isoforms are upregulated following spliceosome component depletion **A)** Boxplots of the log2(fold change) of the MANE (teal) and non-canonical isoforms (dark green) of genes that undergo significant alternative splicing following depletion of the indicated proteins. **B)** Comparison of log2(fold change) of MANE (purple), NMD-targeted (orange), and stable non-canonical isoforms (green) following spliceosome component knockdown. The fold-changes were recalculated using kallisto and DESeq2 after including novel isoforms in the reference transcriptome. Median and number of observations in each group is noted as in Fig 2. P-value above the boxplots is the result of a Wilcoxon test comparing the MANE (purple) and stable non-canonical (green) isoforms to the PTC+ isoforms, with the alternative hypothesis being that the PTC+ isoforms will be more abundant.

It is possible that our ability to accurately quantify levels of annotated transcripts upon spliceosome component knockdown is affected by generation of novel mis-spliced transcripts. These mis-spliced transcripts are not included in the reference sequences used by kallisto for transcript quantification and can affect assignment of sequencing reads to annotated transcripts, thereby altering transcript quantification. To address this issue, we used Stringtie to identify such novel transcripts to include them in the reference list for kallisto-based transcript quantification (**Fig S1**). We found that after accounting for novel transcripts produced upon spliceosome disruption, both non-canonical as well as NMD isoforms show an upregulation as compared to MANE select isoforms in several catalytic spliceosome component depletion datasets (**Fig 4B** and **S5A**). Notably, for a subset of spliceosome component knockdowns (RBM22, EFTUD2, SNRNP200) there is no or only a minor difference in the median fold change values for PTC+ versus other non-canonical transcripts. Thus, we conclude that for this set of spliceosome component knockdowns, a simple comparison of levels of canonical MANE transcripts versus non-canonical transcripts is not sufficient to distinguish the effects of altered splicing from the impact of compromised NMD on the increased abundance of PTC+ transcripts. However, in the case of AQR, SF3B1, SF3B3 and CDC40 depletion, the median fold changes of other non-canonical isoforms are significantly lower than PTC+ isoforms (**Fig 4B**). These conditions are at least somewhat comparable to UPF1, EIF4A3 and MAGOH knockdowns, where levels of canonical and non-canonical isoforms are comparable and significantly lower than PTC+ isoforms. Therefore, it is possible that AQR, SF3B1, SF3B3 and CDC40 have a more direct effect on suppression of PTC+ transcripts by NMD. Still, the increased abundance of non-canonical isoforms as compared to canonical MANE isoforms in these knockdowns suggests that altered splicing could also contribute to the increased levels of PTC+ transcripts.

### Depletion of some catalytic spliceosome components leads to production of novel NMD targeted isoforms in excess of the endogenous NMD targets

Even though our analysis shows that the increased abundance of NMD targets upon depletion of many catalytic spliceosome components cannot be completely attributed to disruption of NMD, the clear increase in levels of PTC+ transcripts under these conditions warrants an investigation for the possible underlying causes. Other groups have previously speculated that disruption of the spliceosome may result in an overabundance of NMD targets, much more than the pathway can handle, thereby overwhelming the NMD machinery and lowering its ability to suppress natural NMD targeted transcripts [41,42]. To test this possibility, we compared the overall levels of annotated and novel transcripts that are targeted to NMD in the spliceosome component versus the control knockdown conditions. An expectation is that in conditions where NMD is overwhelmed, the concentration of novel NMD targets would surpass that of natural NMD targets. We used the Isoform Switch Analyzer algorithm to classify the novel transcripts in the knockdown samples as NMD targets if they contained a PTC more than 50 nt upstream of an exon-exon junction; novel transcripts without a PTC, or with a PTC in the last exon, were classified as stable transcripts [43]. To compare the relative concentrations of annotated and novel transcripts in each sample, we summed the TPM values of all transcripts in each of these groups – canonical MANE, annotated NMD, novel NMD and novel stable. As expected, the MANE isoforms have the highest cumulative TPM estimate in all samples whereas the amounts of annotated NMD isoforms are only fractional (**Fig 5A**, left). Intriguingly, the cumulative amounts of novel NMD transcripts are higher than novel stable transcripts in the case of AQR, SF3B1, SF3B3 and CDC40 knockdowns (**Fig 5A**, right). Curiously, in these four depletion conditions, novel NMD transcripts are almost two-fold or more abundant than annotated NMD transcripts (**Fig 5A**, right inset and **S5B**). It is noteworthy that these four conditions also show a significant increase in PTC+ transcripts as compared to other non-canonical transcripts in **Fig 4B**. Indeed, there is a strong correlation between how significantly the knockdowns upregulate the PTC+ targets and the ratio of total abundance of the novel versus annotated NMD targets (**Fig 5B**). While these results hint at a possibility that excessive production of novel NMD targets could interfere with the suppression of the endogenous NMD substrates, it remains to be seen if a mere doubling of substrate concentration, as is observed in some spliceosome depletion conditions, would be sufficient to bog down the pathway.

**Fig 5.**
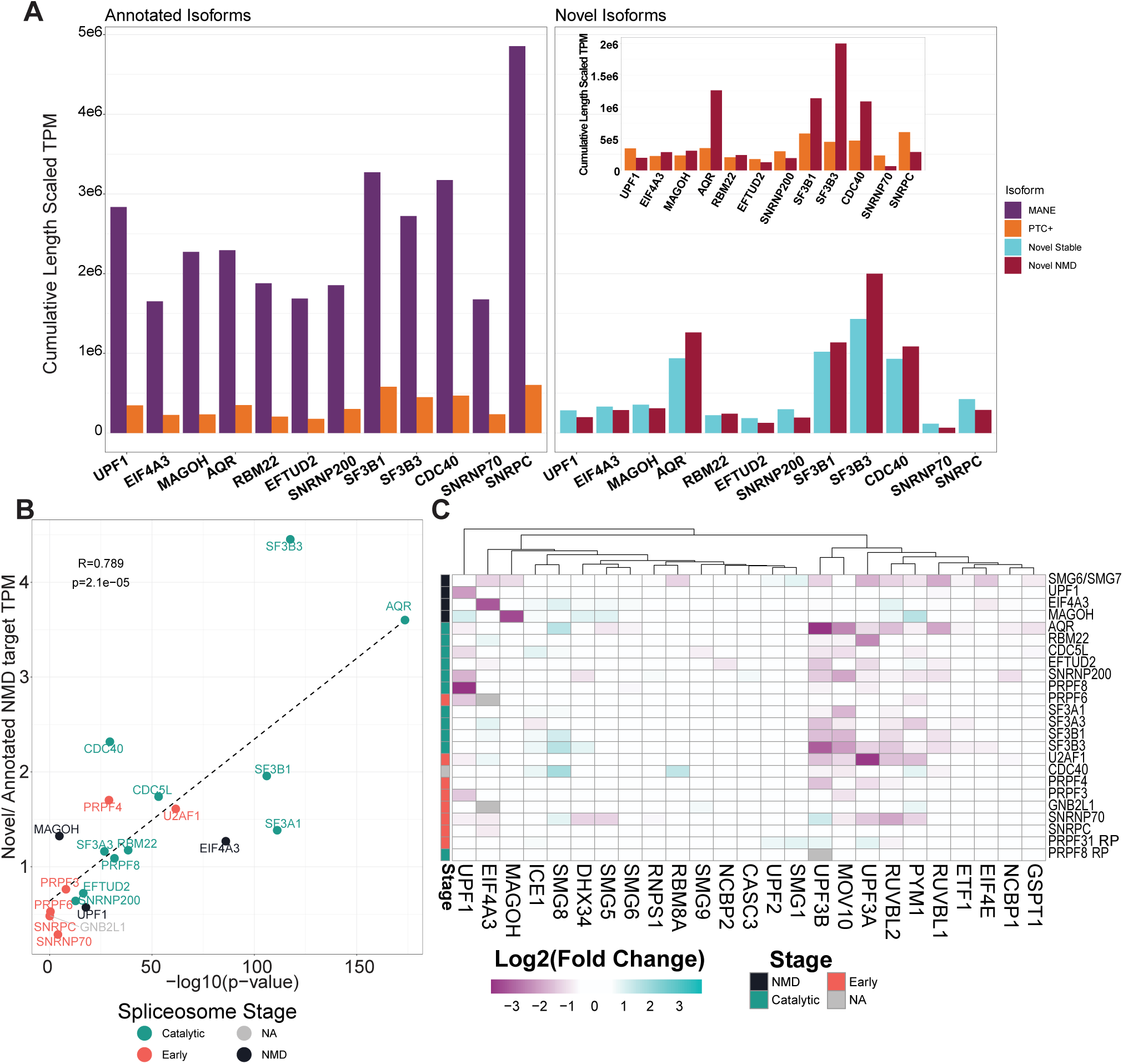
Effect of spliceosome component depletion on relative levels of novel and annotated NMD targeted transcripts and of NMD factor mRNAs. **A)** Left: Cumulative length scaled TPM of MANE (purple) and NMD (orange) transcripts from genes that produce annotated NMD-targeted isoforms in the indicated samples. Right: Cumulative TPMs of stable (blue) and predicted NMD-targeted (red) novel isoforms produced from all genes in the indicated samples. Inset: the cumulative length scaled TPM of annotated NMD-targeted (orange) and novel NMD-targeted transcripts (red), from the left and right plots, respectively, are re-plotted for comparison. In all cases, TPM for each transcript was averaged across replicates before summation. **B)** A scatterplot comparing the ratio of cumulative TPMs of novel:annotated NMD-targeted transcripts (y-axis) to the-log10(p-value) of the upregulation of annotated NMD-targeted transcripts when compared to their MANE counterparts. Dashed line is the linear regression fit with Pearson’s R and p-value shown in upper left corner. **C)** A heatmap clustered along the x-axis showing the log2(fold change) of the MANE isoform of NMD factor genes (x-axis) in spliceosome component or NMD/EJC factor depletion datasets (y-axis). Column labeled “stage” on the left indicates the stage where a spliceosome component leaves the spliceosome as indicated in the legend on the right. Changes less than 1.5 fold in either direction are white, upregulated transcripts are teal, and downregulated transcripts are purple.

### Levels of some NMD factor mRNAs are reduced upon spliceosome disruption

Another possible explanation for the increase of NMD-targeted transcripts upon spliceosome component disruption would be a decrease in abundance of NMD proteins themselves due to altered splicing of their mRNAs thereby causing a partial inhibition of the pathway. To test this possibility, we examined the change in abundance of MANE isoforms of a set of genes that contribute to the NMD pathway [44]. As seen in **Fig 5C**, although MANE isoforms for many NMD factors do not change dramatically upon spliceosome knockdown, there are small clusters of NMD factors whose MANE isoforms are down-or up-regulated in these datasets. The most prominently downregulated set contains the two *UPF3* paralogs, *UPF3A* and *UPF3B*, which are downregulated > 2-fold in multiple catalytic spliceosome component knockdowns tested (4/10 for *UPF3A* and 6/10 for *UPF3B*). Interestingly, this is the same group of spliceosome components whose knockdown leads to an increase in abundance of the NMD-targeted transcripts (**Fig 2**). Similarly downregulated in 6/10 catalytic spliceosome knockdowns (or 8/18 spliceosome factor knockdowns) is the mRNA encoding MOV10. As UPF3A and UPF3B activate both EJC-dependent as well as EJC-independent NMD [39,45], and MOV10 has been suggested to assist UPF1-dependent steps of NMD on some targets [46], downregulation of protein-coding mRNAs of these NMD factors could be a contributor to reduced NMD upon spliceosome disruption. Interestingly, the MANE isoform of SMG8, a regulator of SMG1 kinase, shows a > 2-fold increase in abundance in 4/10 catalytic spliceosome factor depletions. It is notable that while many NMD factor encoding mRNAs are known to be autoregulated by NMD [47,48], we do not observe an upregulation of transcripts encoding NMD factors with the exception of SMG8 and a few other isolated cases. Even upon strong NMD inhibition upon dual depletion of SMG6 and SMG7 proteins in HEK293 cells, only SMG1 and UPF2 encoding MANE isoforms show a > 1.5-fold upregulation. Overall, we conclude that NMD inhibition in K562 cells depleted of spliceosome components could partially result from reduced levels of key NMD activators like UPF3 paralogs and MOV10.

### Spliceosome inhibitor treatment also leads to increased relative abundance of PTC+ and other non-canonical isoforms

Increased abundance of NMD (and in some cases other non-canonical) isoforms in a wide range of spliceosome depletion conditions raises the question if the altered abundance is due to a general effect of spliceosome inhibition rather than a compromised NMD specific function of an individual factor. We argued that this idea can be tested by examining changes in the levels of PTC+, canonical and non-canonical transcripts in human cells treated with spliceosome inhibitors. We performed isoform level quantification of the RNA-Seq data from Naro et al. [49] where prostate cancer 22Rv1 cell line was treated with pladienolide B (an SF3B1 inhibitor [50]), indisulam (targets RBM39 for proteasomal degradation, [51]) and THZ531 (CDK12/13 inhibitor, [52]). Interestingly, similar to the trends we observed with several catalytic spliceosome factor depletion including that of SF3B1, pladienolide B treatment leads to a significant increase in the relative abundance of PTC+ and other non-canonical isoforms as compared to the levels of MANE transcripts from the corresponding genes (**Fig 6A**). Similar trends are observed for indisulam and THZ531 but the median fold changes for PTC+ and non-canonical isoforms are more modest. Notably, RBM39 is among the top 200 targets in one of the four NMD factor genetic screens [26] and it’s indisulam mediated degradation likely resembles individual spliceosome component knockdown. We also compared levels of MANE, PTC+ and other non-canonical transcripts in lymphoblastoid cells treated with high doses of risdiplam [53], which does not inhibit spliceosome but leads to widespread alteration in pre-mRNA splicing by promoting weak splice-site usage [54]. We find that risdiplam treatment also causes an upregulation of NMD targeted transcripts as compared to MANE transcripts although the effect on other non-canonical isoforms is only modest (**Fig 6A**). Notably, treatment with these spliceosome inhibitors and spliceosome activity altering drugs also affects expression of MANE transcripts of various spliceosome components (**Fig 6B**, left) and NMD factors (**Fig 6B**, right). In particular, pladienolide B treatment, which shows highest increase in fold-change of PTC+ and non-canonical isoforms, also exhibits strongest downregulation of mRNAs encoding numerous splicing and NMD factors tested (**Fig 6B**). These observations further indicate that upregulation of NMD transcripts upon spliceosome inhibition, either via chemical inhibitors or individual factor depletion, is likely due to multiple contributing factors that may include reduction in levels of NMD and splicing factors, overproduction of novel NMD substrates and/or dramatically altered gene expression.

**Fig 6.**
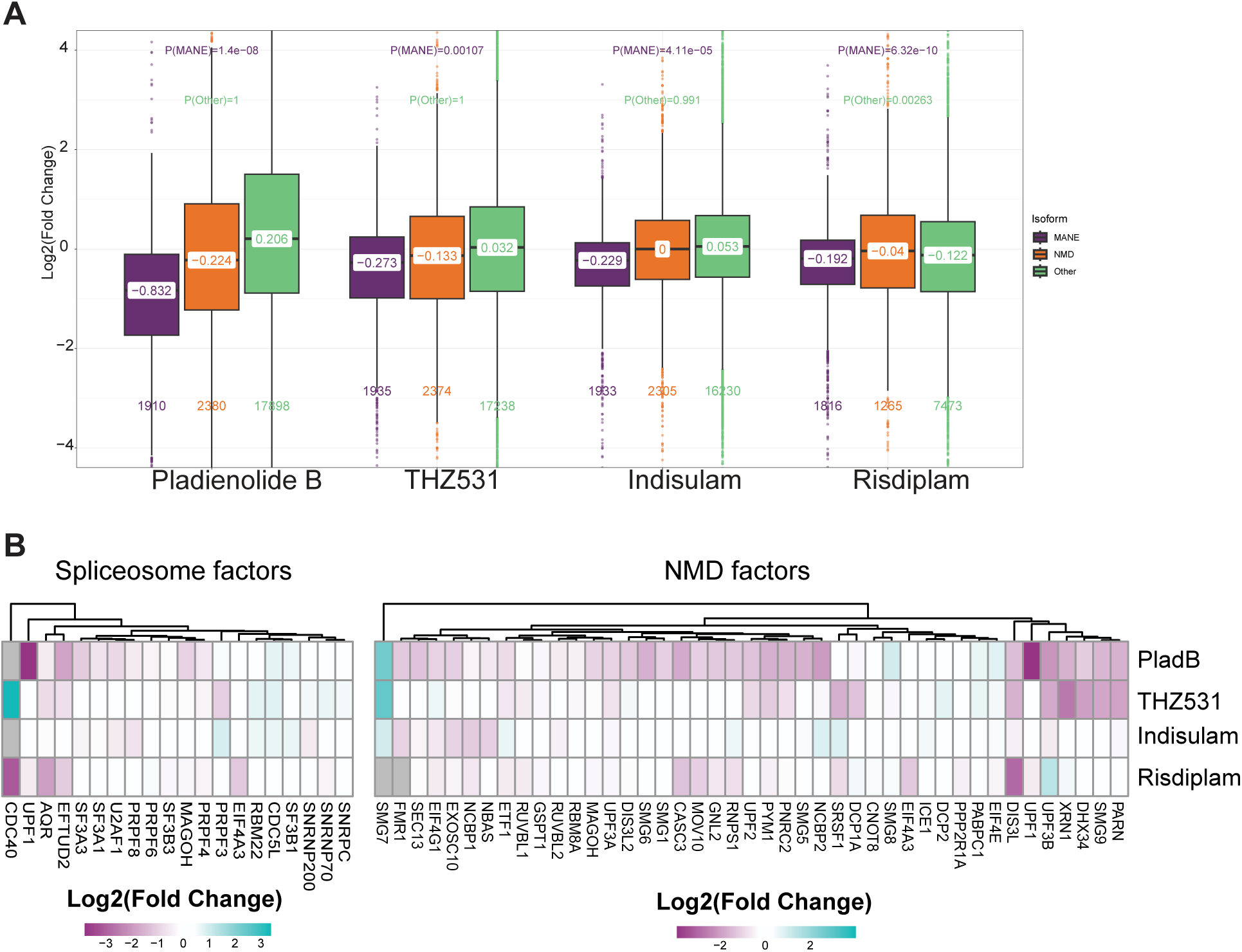
Effects of chemical inhibitors and modulators of spliceosome activity on NMD and other transcripts. **A)** Boxplots showing log2(fold change) of MANE (purple), NMD-targeted (orange), and stable non-canonical isoforms (green) following treatment with splice altering drugs indicated on the x-axis. Medians, numbers of observations and p-values of comparisons shown are as in Fig 4. **B)** Left: A heatmap of the MANE isoform of spliceosome components under investigation (x-axis) following treatment with splice altering drugs (y-axis). Heatmap colors are as in Fig 5. Right: A heatmap of log2(fold change) of MANE isoforms of NMD factors (x-axis) following treatment of splice altering drugs (y-axis). Colors are as in Fig 5.

### Altered gene expression due to disease causing mutations in spliceosome components also includes increased abundance of NMD targeted isoforms

Based on our findings above, we predict that spliceosome mutations that cause human disorders will result in increased abundance of NMD targets in addition to altering pre-mRNA splicing. We investigated this possibility by examining available RNA-Seq datasets from retinitis pigmentosa (RP) patient-derived fibroblasts with a mutation in *PRPF8* and induced pluripotent stem cells (iPSCs) made from patient-derived fibroblasts with a mutation in *PRPF31*. The *PRPF8* deficient fibroblasts were derived from a patient with a deletion *(c6974-6994del)* that disrupts a region required for interaction with EFTUD2 and SNRNP200 [55] whereas the iPSCs with *PRPF31* deficiency were from a patient with very severe RP caused by deletion (*c1115-1125del11*) that causes a frameshift leading to a truncated protein [56]. When compared with cells derived from the control normal individuals, cells with either *PRPF8* or *PRPF31* mutation show an upregulation of PTC+ (**Fig 7A**) or poison exon-containing transcripts **(Fig 7B)** as compared to their MANE counterparts. In the case of *PRPF31* mutant cells, the upregulation of PTC+ appears to be somewhat more specific and significantly higher than for non-canonical isoforms. Notably, the effect of mutations in PRPF8 are similar to what we observe from *PRPF8* knockout K562 cells from ENCODE (**Fig S2A**). As a further confirmation of increased abundance of NMD targets in these cells, read distribution across *EIF4A2* gene locus shows increased inclusion of an NMD-targeting poison exon in the cells with mutant *PRPF31* mutant cells, though in cells with mutant *PRPF8* there is little difference from the wildtype sample (**Fig 7C**).

**Fig 7.**
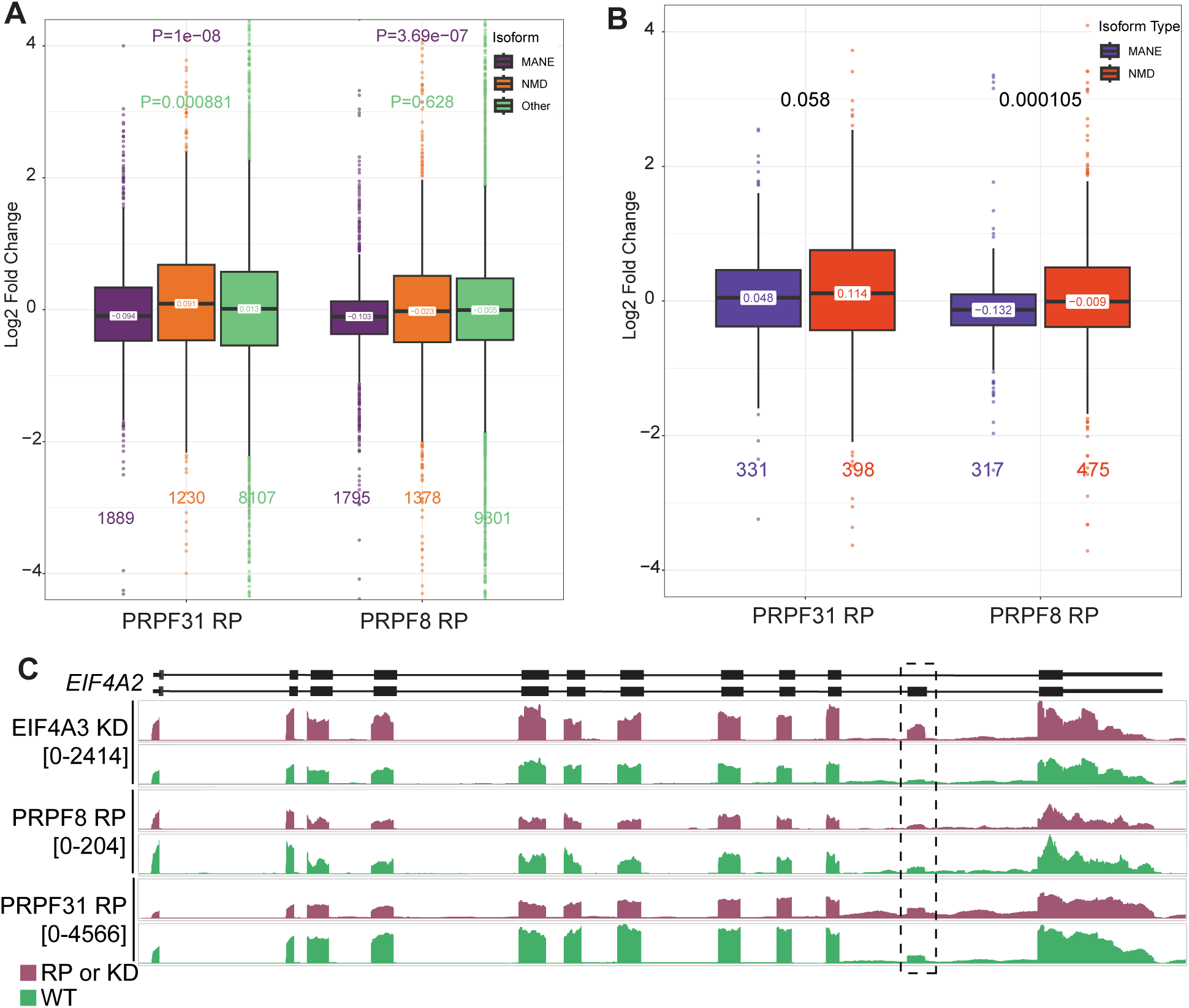
Effects of retinitis pigmentosa causing mutations in *PRPF8* and *PRPF31* on NMD targeted and other isoforms in patient-derived cells. **B)** Boxplots showing log2(fold change) distributions of MANE (purple), NMD-targeted (orange), and stable non-canonical isoforms (green) in cells with retinitis pigmentosa causing mutations in *PRPF31* or *PRPF8* (x-axis). Medians, numbers of observations and p-values of comparisons shown are as in Fig 4. **C)** Boxplots as in A showing log2(fold change) distributions of MANE (blue) and NMD-targeted isoforms (red) of genes containing poison exons in cells with retinitis pigmentosa causing mutations in *PRPF31* or *PRPF8*. **D)** RNA-Seq read distribution on *EIF4A2* gene locus in representative replicates of cells derived from retinitis pigmentosa patients (purple) or normal individuals (green). The poison exon in *EIF4A2* is indicated by the dashed box. The scale of each pair of tracks is indicated under the sample name.

Like in the case of several catalytic spliceosome component knockdowns, iPSCs with the *PRPF31* mutation show a reduced abundance of MANE isoforms from genes that undergo alternative splicing (**Fig S6**). However, the same pattern is not seen in patient fibroblasts with mutations in *PRPF8*. Thus, *PRPF31* and *PRPF8* mutations could also alter expression of key NMD regulated genes that may contribute to disease phenotypes. One such example is *NRG1*, which plays a role in motor and sensory neuron development. In RP patient-derived *PRPF8* mutant cells, the NMD-targeted isoform of *NRG1* has a log2 fold change of 1.34 (padj = 0.001, **Table S5**) while the MANE isoform only changes mildly. Though RP is not a cranio-facial disorder, NMD-targeted isoforms of genes that are involved in cranio-facial development are also upregulated in *PRPF31* mutant cells, hinting at shared gene expression mechanism underlying different types of spliceosomopathies. One example is *PDLIM7*, which encodes a scaffold protein that localizes LIM-binding proteins to actin filaments and is involved in formation of bones including flat bones in mandible and cranium [57]. In RP patient-derived *PRPF31* mutant cells, a poison exon-containing NMD isoform of *PDLIM7* is upregulated (log2FC 1.14, padj = 0.04; **Table S5**) while its protein-coding MANE isoform is mildly downregulated. Notably, similar changes in *PDLIM7* PTC+ and MANE isoforms are also observed in *PRPF8* mutant cells. Other NMD regulated isoforms upregulated in *PRPF8* and *PRPF31* mutant cells include well-known NMD targets such as *SRSF2* and *SRSF3* as well as isoforms of genes with functions that may contribute to disease progression [e.g., *OSTM1* (regulates chloride channels in osteoclasts and melanocytes)]. These observations indicate that even if the effect of spliceosome factor mutations on NMD target abundance could be pleiotropic in nature, disruption of some key NMD-regulated activities potentially contribute to disease progression in spliceosomopathies.

## DISCUSSION

### Spliceosome disruption increases abundance of NMD substrates

The identification of a surprisingly large number of spliceosome components in several genetic screens for NMD factors in human cell lines indicates that our understanding of the impact of spliceosome function on NMD remains incomplete. In particular, proteins present in the spliceosome when it performs the two catalytic steps of splicing appear to be more influential on NMD (**Fig 1**). Motivated by these observations, our further investigation has revealed that, indeed, when spliceosome function is disrupted due to depletion of one of its many core components, transcripts normally suppressed by the NMD pathway are upregulated (**Fig 2**). Our analysis confirms the broad impact of the spliceosome core on NMD as the upregulation of PTC+ transcripts is observed in 15/18 spliceosome proteins tested. Moreover, the increased abundance of NMD-targeted isoforms is also seen in cells treated with spliceosome inhibitors (**Fig 6**) and in the transcriptomes of cells derived from human patients with spliceosome component mutations (**Fig 7**). As expected, depletion of spliceosome core proteins leads to widespread changes in pre-mRNA splicing (**Fig 3**), which alters overall gene expression (**Fig 3** and **4**). Even after we tune transcript quantification to account for these alterations in the transcriptome, depletion of the spliceosome components shows increased abundance of EJC-dependent NMD targets as a group as compared to the canonical transcripts (**Fig 4B and S5A**), indicating that NMD dependent gene regulation is compromised when the spliceosome is dysfunctional.

The effect on NMD substrate abundance is more pronounced upon reduction of catalytic spliceosome components as compared to U1 and U4 components that function during early steps. (**Fig 2, S2 and S5A**). Depletion of all 11 catalytic spliceosome components causes a highly significant upregulation of PTC-containing transcripts whereas in the case of early components either the effect on NMD substrates is insignificant (SNRNPC and PRPF6) or only moderately significant (SNRNP70 and PRPF3) (**Fig 2 and S2).** Among the factors that are not part of the catalytic spliceosome, depletion of only U2AF1 and PRPF4 leads to a strong effect on NMD targets. A more consequential effect of catalytic spliceosome components on NMD could be due to compromised EJC assembly (see below). We notice that, as compared to the catalytic components, early spliceosome factor depletion affects a fewer number of annotated splicing events (**Fig 3A**) and a weaker effect on the levels of NMD factor mRNAs (**Fig 5C**). Intriguingly, a recent study shows that in HEK293 cell lines depletion of U1 components results in more splicing changes than that of catalytic spliceosome components [58]. One possibility is that the increase in altered splicing events from early spliceosome component depletion could hinder K562 cell survival. If this were the case, the early spliceosome components would be less amenable to acute depletion experiments, possibly explaining their under-representation in the ENCODE database and in screens for NMD factors (**Fig 1**). If depletion of many early spliceosome components is lethal to cells, it stands to reason that the only early components that survive acute protein depletion are the ones with a milder effect on gene expression including NMD.

### Possible causes of NMD inhibition upon spliceosome disruption

The NMD defects upon spliceosome disruption could result from either a direct interference of the pathway or due to indirect effects. The direct effects could result from compromised EJC assembly, which is initiated upon recruitment of EIF4A3 by the CWC22/CWC27 heterodimer to the B^act^ spliceosomal complex [14–17,59]. EIF4A3 and/or the assembled pre-EJC bound to a contiguous stretch of 6 nt in the 5’-exon is observed in the C complex [60,61]. While CWC22 and CWC27 depletion datasets were not available in the ENCODE database for us to test their effects on global NMD, knockdown of CWC22 in HeLa cells has been previously shown to upregulate levels of NMD targets [14]. Thus, reduced ability to recruit/deposit EJC proteins on exonic RNA within the spliceosome can directly impact NMD. Another candidate for a role in EIF4A3 recruitment is EFTUD2, which sits adjacent to EIF4A3 in the C complex where its C-terminus engages with the RecA domain of EIF4A3 [60]. Intriguingly, like EIF4A3, mutations in EFTUD2 cause a disorder characterized by cranio-facial defects and intellectual disability [22,23,62,63]. Although EFTUD2 knockdown upregulates PTC+ transcripts, similar effects are also observed for non-NMD targets (**Fig 4B**). Thus, in this case, we cannot separate direct effects of EFTUD2 on EJC/NMD from its possible indirect effects (see below).

We identified four components of the catalytic spliceosome that have a specific effect on PTC+ isoforms (**Fig 2 and 4B**). First among these is AQR, which has been previously shown to contribute to EJC deposition and NMD [18]. Indeed, we observe that AQR depletion in K562 cells causes a strong and specific upregulation of PTC+ isoforms as compared to both MANE as well as other non-canonical isoforms (**Fig 2 and 4B**). A similar effect on NMD targeted isoforms is observed in the case of two components of the SF3B complex of the U2 snRNP, SF3B1 and SF3B3 (**Fig 2 and 4B**). Notably, Cheruiyot et al, recently showed that SF3B1 and U2AF1 variants carrying myelodysplastic syndrome-causing mutations cause upregulation of NMD-targeted endogenous as well as reporter mRNAs in K562 cells [28]. The effects of SF3B1 inhibition on NMD are also supported by the upregulation of NMD substrates by SF3B1 targeting spliceosome inhibitor, pladienolide B (**Fig 6**). Possibly, SF3 complexes within the U2 snRNP may also have a role in determining the potential of a spliced RNA to undergo NMD as several other components of SF3 complexes are strongly enriched in NMD factor screens (**Table S3**). Our results from SF3B3 knockdown K562 cells support such a possibility (**Fig 2 and 4B**). It is interesting to note that in the catalytic spliceosome SF3A and SF3B complexes and AQR bind adjacent to the intron in a region close to the branchpoint. Moreover, the intron binding complex nucleated by AQR can be chemically crosslinked to SF3A and SF3B proteins [64]. Lastly, another protein that has a specific effect on PTC+ isoforms is CDC40 (also known as Prp17), a step II factor (**Fig 2** and **4B**) [32]. CDC40 interacts with multiple protein and RNA components within the catalytic spliceosome including the U2-branch site helix, U6 snRNA and U5 proteins. Through these interactions, it plays a crucial role in stabilizing the second-step conformation of the spliceosome [60,61]. A hypothesis that emerges from these observations is that protein components that properly position the intron within the catalytic spliceosome also directly impact recruitment/deposition of EJC subunits and thereby reduce the potential of a spliced mRNA to be regulated by NMD.

In addition to the possible direct effects on NMD through EJC deposition, disruption of spliceosome function by reduced levels or mutations in its core components also causes pleiotropic effects that contribute to impaired NMD. One indirect effect could be due to mis-regulation of genes encoding NMD factors such as the two UPF3 paralogs, which enhance both EJC-dependent and EJC-independent NMD [65,66]. Although mRNAs encoding UPF3 factors are particularly sensitive to spliceosome disruption (**Fig 5C**), it remains to be seen if their protein levels are also reduced. We also examined a previously proposed hypothesis that overproduction of novel NMD substrates due to mis-splicing could indirectly affect the ability of the pathway to regulate its normal targets [65,66]. Interestingly, in cells depleted of the four factors that produce a significant and specific effect on NMD targeted isoforms, we observe that overall concentration of novel NMD substrates is at least 2-fold more as compared to annotated NMD targets (**Fig 5A**). In various knockdowns tested, we even observe a significant direct correlation between the relative amounts of novel NMD transcripts detected and how significant the NMD target upregulation is in each knockdown (**Fig 5B**). Thus, the amount of novel NMD targets generated could influence the degree of NMD inhibition. It remains to be seen if a mere two-fold increase in NMD target concentration is sufficient to overburden the pathway. Such an outcome seems less likely considering a previous report that NMD activity is stable across tissues [67], where concentration of NMD substrates is expected to be highly variable. Indirect effects on NMD targets could also stem from a departure from normal splicing patterns and consequential increase in production of non-canonical transcripts including NMD isoforms. This possibility is supported by the reduced abundance of canonical isoforms and increased levels of non-canonical transcripts that are not targeted to NMD but are transcribed from the same set of genes in several knockdowns (**Fig 4B**), spliceosome inhibitor treated cells (**Fig 6**) and RP patient-derived cell lines (**Fig 7**). Experimental strategies that can differentiate RNA production and degradation rates will be necessary to parse out contributions of such indirect effects on NMD target abundance. Finally, due to the cross-regulation between the splicing machinery [58], inhibition of splicing will also alter spliceosome factor abundance to further compound these indirect effects. Most likely, the increased abundance of endogenous NMD targets observed in the analysis presented here and of NMD reporter RNAs in the recent genetic screens [26–29] results from a combination of direct and indirect effects. Direct effects may be limited to a smaller set of spliceosome components whereas reduced/lost function of most spliceosome factors is expected to result in at least some pleiotropic effects. Regardless of the exact mechanism, our observations support a conclusion that compromised NMD based regulation is another hallmark of cells with impaired spliceosome function.

### Consequences of NMD substrate misregulation in spliceosomopathies

Many genes that are important for developmental processes and cell differentiation pathways rely on NMD for their regulation [68–70]. A prominent group comprises genes containing poison exons, which are enriched in developmental functions [6]. Our results show that NMD targeted isoforms generated from this set of genes are upregulated upon spliceosome disruption including in cells derived from two patients with retinitis pigmentosa mutations in *PRPF8* and *PRPF31* (**Fig 7**). Among these are developmentally important genes related to neuronal growth (e.g. *NRG1*, **Table S5**). Interestingly, in these RP datasets we also observe upregulation of some genes with functions in cranio-facial development (e.g., *PDLIM7* and *OSTM1,* **Table S5**). These findings, in conjunction with the increased abundance of NMD targeted mRNAs upon depletion of other spliceosome components that cause human disorders when mutated (e.g., EFTUD2, SNRNP200, PRPF4, PRP17, SF3B1), suggest that impaired NMD is likely to be a contributing factor in these disorders. In conclusion, although the exact cause of NMD target upregulation upon spliceosome disruption remains to be determined, we recommend that future investigations into spliceosomopathies should consider NMD disruption as a likely contributor to their molecular phenotypes.

## METHODS

### Identification of splicing factors in NMD screens

Extended data tables of hits in NMD screens were downloaded from previous studies [26–29]. To identify high confidence hits we restricted each list to the top 200 factors based on the rankings in the original studies. Using custom R scripts, we determined the overlap between all four lists. A complete list of human spliceosome components were downloaded from the spliceosome database [31], and the list of potential NMD factors were annotated based on a gene’s presence in the spliceosome database. The last complex of the spliceosome a protein is associated with was determined by selecting the last formed complex listed for that protein in the spliceosome database. STRING and GO biological process analysis, with filtering for redundant terms set at 0.75, was conducted using Cytoscape version 3.10.1 [74].

### RNA-Seq datasets and their processing

RNA-seq datasets from siRNA knockdown or CRIPSR-Cas mediated knockout of spliceosome components identified in 2 or more of the NMD screens, as well as other spliceosome components of the U1, U2, U4, U5, and U6 components were identified from the ENCODE consortium’s ENCORE project [33–35] and retrieved from the SRA archive (**Table S1**). Reads were trimmed using trimmomatic version 0.36 [75] to remove adaptors, bases with a quality less than 3, and reads shorter than 30nt. Further quality control and analyses performed on these datasets are visually represented in **Fig S1**.

### Generation of NMD target lists

Transcripts containing a predicted PTC were previously described [39]. To obtain a list of NMD targets, we analyzed the SMG6 knockdown and SMG7 combined knockout from Gehring and colleagues, and identified transcripts that are more than 1.5 fold upregulated [12]. Any PTC-containing transcripts that are upregulated > 1.5 fold in SMG6 and SMG7 double-depletion were considered as NMD targeted PTC+ transcripts. BioMart was used to retrieve the isoform characteristics from the ENSEMBL GRch38 build 109 of the human genome, including MANE select transcript status and transcript biotype from each isoform on the PTC+ list [76]. Genes where the MANE select transcript was on either the PTC- transcript or the NMD-transcript list were removed from the dataset. MANE transcripts from this set of genes were used as the MANE isoform group, and non-MANE non-NMD biotype isoforms were used as the stable non-NMD transcript group.

The poison exon target list was made by identifying previously reported genes that contain conserved poison exons which introduce a PTC [5]. All transcripts from these genes were obtained via BioMart. Transcripts annotated with the nonsense mediated decay biotype were included as the NMD targets, and the MANE select transcript were included as the non-NMD targets. Genes with a MANE select transcript with a NMD biotype or on the NMD target list were excluded.

### Differential expression analysis

Reads were mapped and quantified using kallisto version 0.43.1, based on an index generated from cDNA from Ensembl GRch38 build 109 [76]. Kallisto was run on paired end reads using default settings [36]. Transcript quantification was imported to R using tximport which calculated length scaled TPM. Transcripts with a mean length scaled TPM less than 1 in wildtype or knockdown samples were filtered out. Principal component analysis (PCA) was conducted in R using the DEseq2 package on the raw counts using default settings, comparing the WT and depletion conditions. ENCODE datasets without clear segregation between treatment conditions were removed from the study (**Fig S1**). Differential expression analysis was conducted comparing KD to WT transcript levels with DESeq2 using default settings [37]. ENCODE datasets where the MANE isoform of the spliceosome component being depleted were not more than 1.5-fold downregulated were also removed from the study (**Fig S1**). Gene level differential expression analysis was conducted as above, however the tx2gene option was specified, and a table of transcript and corresponding gene IDs were provided to convert transcript level counts to gene level.

The results of differential expression analysis were annotated based on the characteristic under investigation for each plot, and the log2fold change of transcripts on those lists were plotted using ggplot2 [77]. A Wilcoxon test was used to determine statistical significance between two groups.

### Splicing analysis

Splicing analysis was performed with rMATS turbo version 4.2.0, [40] using binary index and GTF from ensemble GRch38 build 109. A read length of 100 was specified for datasets from ENCODE, while the average read length from all samples was used for other datasets. The –variable-read-length and –novelSS flags were used to identify novel splicing events created following spliceosome knockdown. Splicing changes were calculated using reads mapping to exon-exon junctions and exons (labeled by rMATS as JCEC). Splicing changes were considered significant if padj < 0.05. Genes were classified as undergoing alternate splicing in a dataset if there was one or more significant splicing changes or novel splicing events. Total number of significant or novel splicing events were normalized according to average number of reads mapped by HISAT2 across all samples from an experiment. Results of the splicing analysis were used to annotate the output of differential expression analysis to compare genes that do or do not experience altered splicing.

### Novel isoform analysis

Following the IsoformSwitchAnalyzeR documentation, novel isoforms were identified by using HISAT2 v2.1.0 for mapping, and Stringtie v1.3.3b for isoform quantification [78,79]. HISAT2 used an index with annotated splice sites and exons built from Ensembl GRch38 build 109 and was run with the-dta options. Following HISAT2, reads were aligned and merged by StringTie using a reference annotation from Ensembl GRch38 build 109. Finally, the reads were quantified using StringTie with the-e option specified, using the merged transcriptome as the reference annotation.Transcripts and quantifications were imported into R and analyzed via IsoformSwitchAnalyzeR v2.2.0 [43]. During the importRdata step the merged transcriptome from StringTie was used as the exon annotation and sequences from the same file extracted via gffread v0.12.7 [80] were used as the fasta file. The default settings were used for pre-filtering and switch testing using DEXseq. Open reading frame (ORF) analysis was added using the same GTF file used for StringTie, and novel ORF analysis was conducted with default settings. The consequence of isoform switching was determined based on NMD status. Transcripts were classified as NMD targets by IsoformSwitchAnalyzeR using the program’s default settings based on the 50nt rule.

The sequence of novel isoforms were extracted with IsoformSwitchAnalyzeR and were used along with the transcriptome from Ensembl GRch38 build 109 to make a kallisto index for each dataset. Kallisto quantification was conducted as described above using the index containing novel transcripts. Transcripts with length scaled TPM less than 1 in both WT and KD datasets were filtered out before the DESeq2 step as above. The length scaled TPM for transcripts calculated by tximport was used to compute and compare cumulative TPM for different classes of transcripts.

### Analysis of non-ENCODE datasets

Datasets from cells treated with Pladienolide B, THZ531, and Induslam [49] were retrieved from the SRA and processed as above. Differential expression analysis was completed as above, with treatment datasets compared to DMSO treated data. RNA-seq datasets from cells treated with risdiplam [53] were retrieved from the SRA database and technical replicates were merged into one dataset. Reads were trimmed and quantified as above. We considered the RNA-seq datasets treated with the two highest concentrations of risdiplam, 3160 and 10000 nM, as two replicates of the treatment condition and compared them to cells treated with DMSO in differential expression analysis. Datasets from iPSC cells from healthy patients and patients with *PRPF31* mutations [56] and from fibroblast cells from healthy patients and patients with *PRPF8* mutations [55] were retrieved from the SRA and quantified as above. Differential expression analysis for all datasets were conducted using DESeq2 as above. SRA IDs of all non-ENCODE datasets are also listed in **Table S1**.

## Data Availability

All code written in support of this publication is publicly available at https://github.com/ceOSU/Embree-et-al-2025-Direct-and-indirect-effects-of-spliceosome-disruption. All datasets examined in this study are listed in **Table S1**.

## AUTHOR CONTRIBUTIONS

Conceptualization: C.M.E. and G.S.; Formal analysis: C.M.E. and A.S.; Investigation: C.M.E. and A.S.; Writing – Original draft and preparation: C.M.E. and G.S.; Writing – Review and editing: C.M.E. and G.S.; Funding acquisition, Project Administration and Supervision: G.S.

## ACKNOWLEDGEMENTS

This work was supported by grants from NIH (R01-GM120209 and R35-GM149298) to G.S. C.M.E. was partially supported by an NIH T32 training grant (T32-GM141955). We also acknowledge an allocation of computational resources from the Ohio Supercomputer Center.

## SUPPORTING INFORMATION CAPTIONS

**Fig S1.** Analysis pipeline for RNA-seq datasets

Flowchart of the experimental pipeline showing the steps of analysis used. Steps where datasets were used from further analysis is indicated in red.

**Fig S2.** Depletion of many catalytic spliceosome components upregulates endogenous NMD targeted mRNAs in K562 cells.

**A)** Boxplots displaying the log2(fold change) on the y-axis of MANE transcripts (purple) and NMD-targeted transcripts (orange) in the knockdowns indicated on x-axis. The number of transcripts in each group is indicated below the boxplot, the median of the boxplot is indicated on the boxplot, and the p-value of the Wilcoxon text comparing the two groups is above. Spliceosome component names are colored according to when they leave the spliceosome as in A.

**B)** Boxplot showing the log2(fold change) of MANE transcripts (blue) and NMD-targeted transcripts (red) from genes with conserved poison exons. Boxplot and depletion annotations are as in B.

**Fig S3**. Widespread changes in annotated and novel splicing events upon spliceosome factor knockdowns reduce expression of the affected genes.

**A)** The log2(fold change) of genes that are (green) and are not (pink) undergoing altered splicing following spliceosome component knockdown. Comparisons are made at the gene level.

**B)** The log2(fold change) of genes that are (green) and are not (pink) undergoing altered splicing following spliceosome component knockdown. Comparisons are made at the MANE transcript level

**Fig S4.** MANE isoforms are downregulated in genes undergoing alternate splicing but not in genes without altered splicing patterns.

**A)** Boxplots of the log2(fold change) of the MANE (pink) and non-canonical isoforms (dark red) of genes that do not undergo significant alternative splicing following depletion of the indicated proteins.

**B)** Boxplots of the log2(fold change) of the MANE (teal) and non-canonical isoforms (dark green) of genes that undergo significant alternative splicing following depletion of the indicated proteins.

**Fig S5.** Effect of spliceosome component depletion on relative levels of novel and annotated NMD targeted transcripts and of NMD factor mRNAs.

**A)** Comparison of log2(fold change) of MANE (purple), NMD-targeted (orange), and stable non-canonical isoforms (green) following spliceosome component knockdown. The fold-changes were recalculated using kallisto and DESeq2 after including novel isoforms in the reference transcriptome. Median and number of observations in each group is noted as in Fig 2. P-value above the boxplots is the result of a Wilcoxon test comparing the MANE (purple) and stable non-canonical (green) isoforms to the PTC+ isoforms, with the alternative hypothesis being that the PTC+ isoforms will be more abundant.

**B)** Left: Cumulative length scaled TPM of MANE (purple) and NMD (orange) transcripts from genes that produce annotated NMD-targeted isoforms in the indicated samples. Right: Cumulative TPMs of stable (blue) and predicted NMD-targeted (red) novel isoforms produced from all genes in the indicated samples.

**Fig S6.** Genes undergoing alternate splicing are downregulated in as a result of some spliceosome component mutations.

The log2(fold change) of genes that are (green) and are not (pink) undergoing altered splicing in cells with retinitis pigmentosa causing mutations in spliceosome components. Comparisons are made at the MANE transcript level

**Table S1.** SRA accession numbers for the RNA-seq datasets analyzed in this study.

**Table S2**. Gene ontology biological processes enrichment chart for the top 200 proteins identified in each NMD screen, and the proteins shared in 2 or more of the screens.

**Table S3**. Splicing factors identified in the top 200 factors in any of the four NMD genetics screens.

**Table S4.** Results of DEseq2 for all transcripts found following spliceosome component depletion by the ENCODE consortium.

**Table S5.** Results of DEseq2 for transcripts on the PTC+ gene list in cells containing retinitis pigmentosa causing mutations in spliceosome components.

