## Supplemental Figure 1 for "Direct and indirect effects of spliceosome disruption compromise gene regulation by Nonsense-Mediated mRNA Decay"

### Figure S1

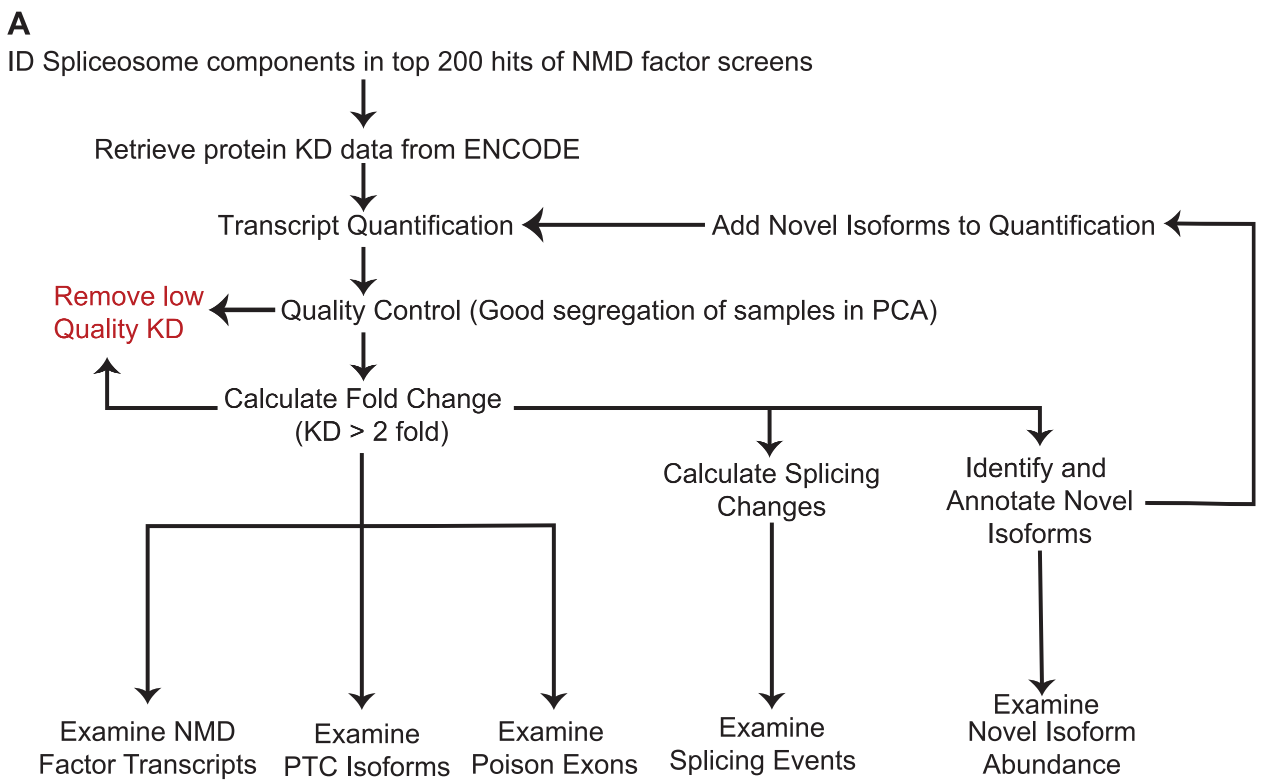


**Fig S1. Analysis pipeline for RNA-seq datasets**
Flowchart of the experimental pipeline showing the steps of analysis used. Steps where datasets were used from further analysis is indicated in red.
