## Supplemental Figure 2 for "Direct and indirect effects of spliceosome disruption compromise gene regulation by Nonsense-Mediated mRNA Decay"

### Figure S2

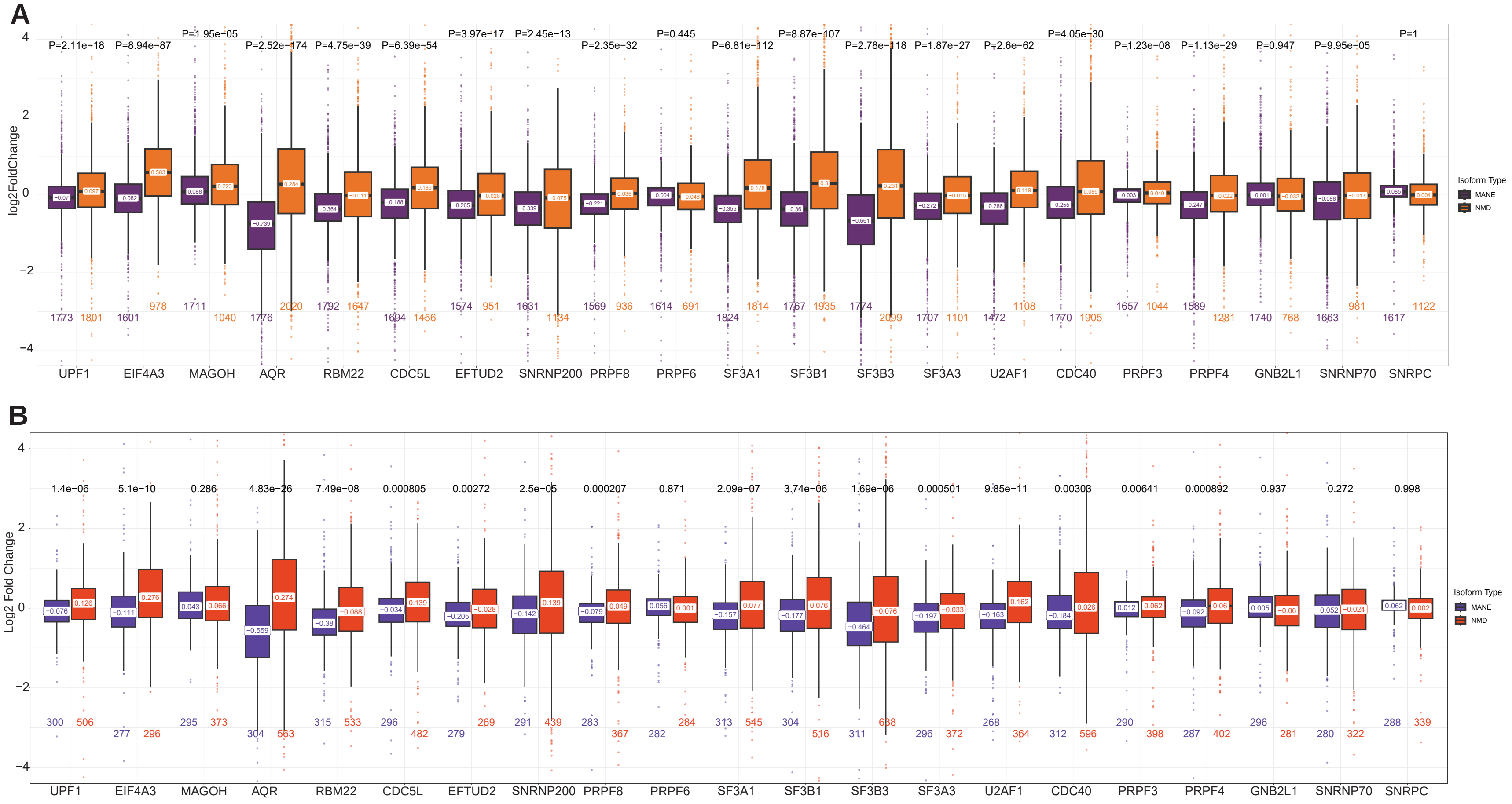


**Fig S2. Depletion of many catalytic spliceosome components upregulates endogenous NMD targeted mRNAs in K562 cells.**
**A)** Boxplots displaying the log2(fold change) on the y-axis of MANE transcripts (purple) and NMD-targeted transcripts (orange) in the knockdowns indicated on x-axis. The number of transcripts in each group is indicated below the boxplot, the median of the boxplot is indicated on the boxplot, and the p-value of the Wilcoxon text comparing the two groups is above. Spliceosome component names are colored according to when they leave the spliceosome as in A.
**B)** Boxplot showing the log2(fold change) of MANE transcripts (blue) and NMD-targeted transcripts (red) from genes with conserved poison exons. Boxplot and depletion annotations are as in B.
