## Supplemental Figure 3 for "Direct and indirect effects of spliceosome disruption compromise gene regulation by Nonsense-Mediated mRNA Decay"

### Figure S3

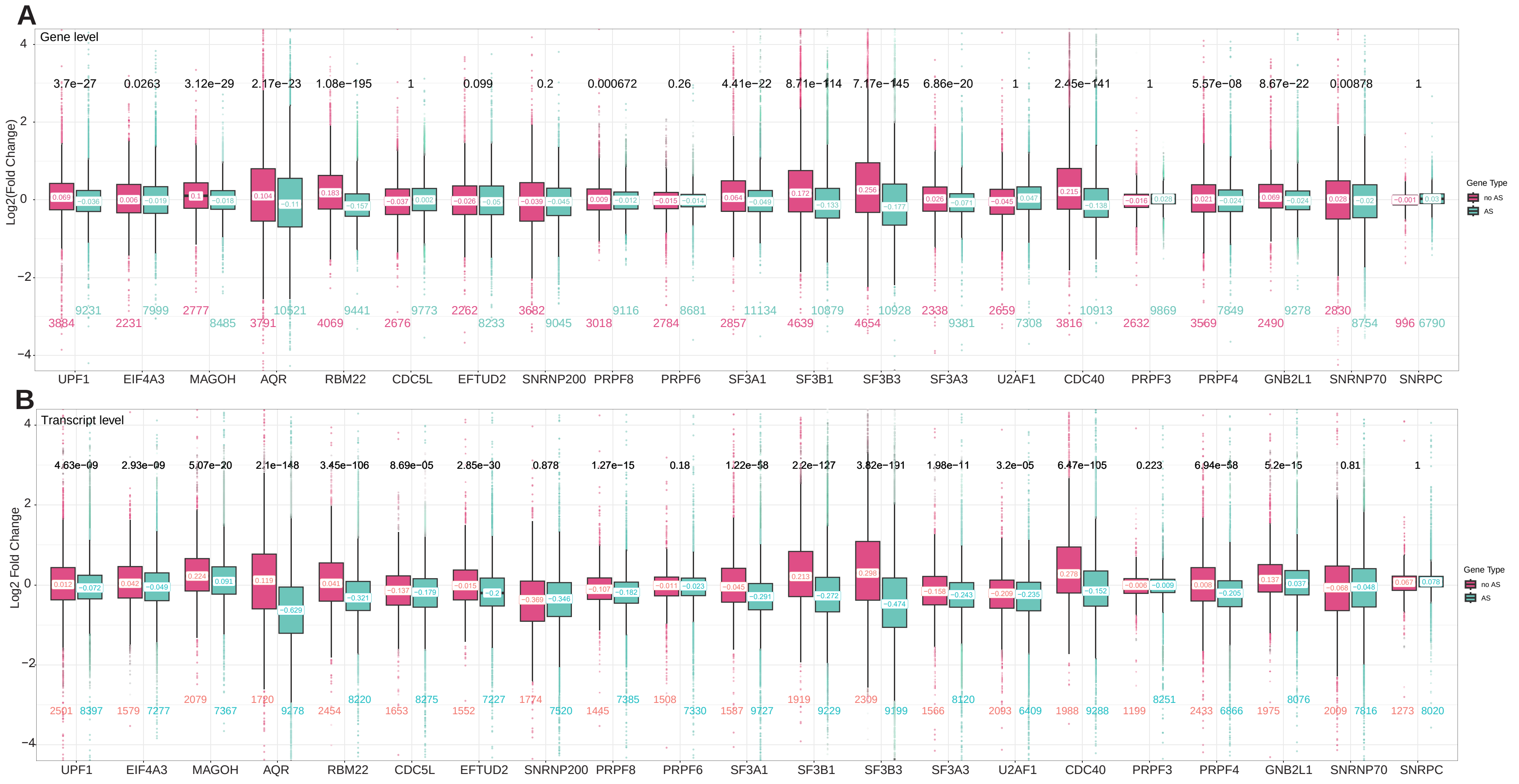


**Fig S3. Widespread changes in annotated and novel splicing events upon spliceosome factor knockdowns reduce expression of the affected genes.**
**A)** The log2(fold change) of genes that are (green) and are not (pink) undergoing altered splicing following spliceosome component knockdown. Comparisons are made at the gene level.
**B)** The log2(fold change) of genes that are (green) and are not (pink) undergoing altered splicing following spliceosome component knockdown. Comparisons are made at the MANE transcript level
