## Supplemental Figure 4 for "Direct and indirect effects of spliceosome disruption compromise gene regulation by Nonsense-Mediated mRNA Decay"

### Figure S4

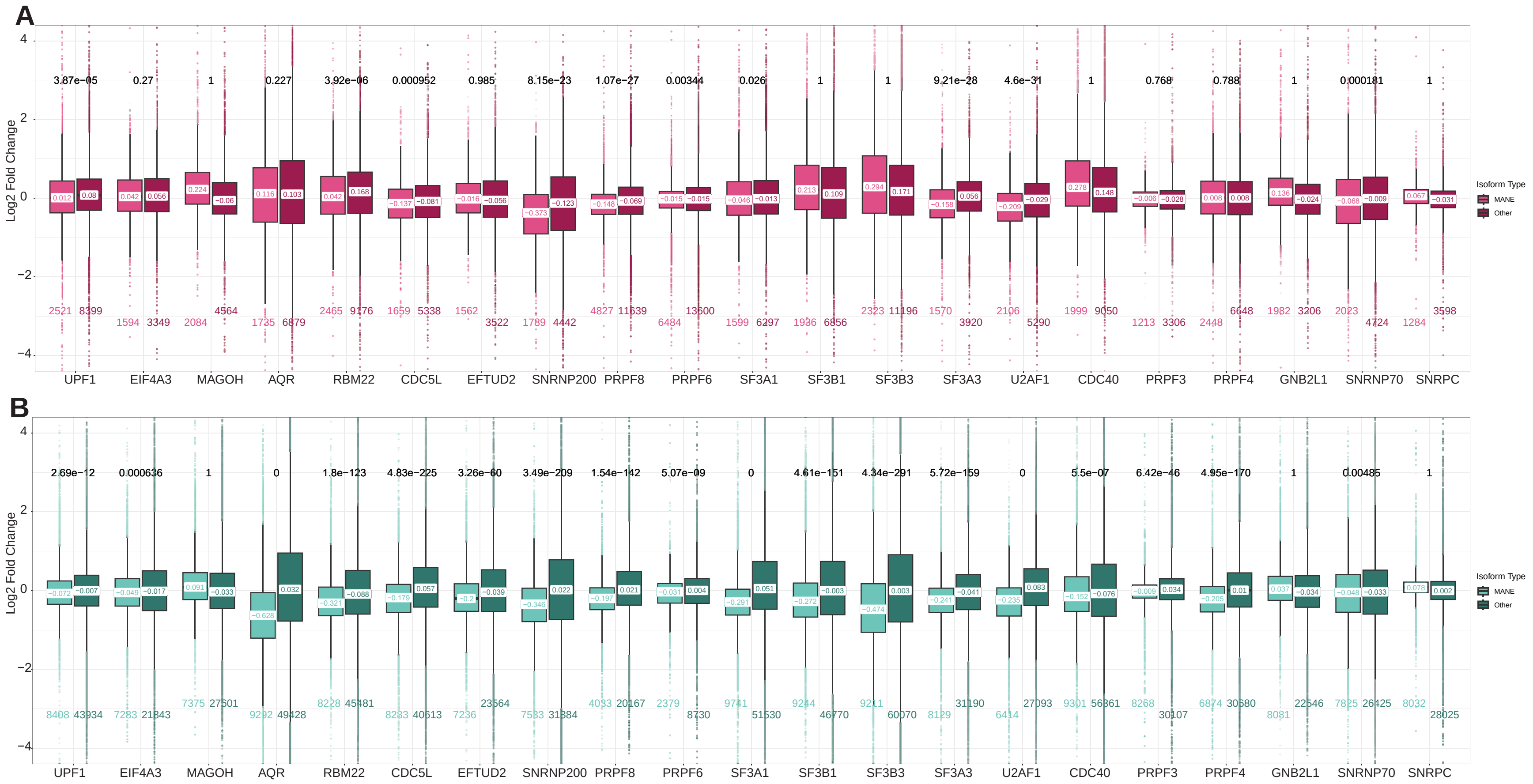


**Fig S4. MANE isoforms are downregulated in genes undergoing alternate splicing but not in genes without altered splicing patterns.**
**A)** Boxplots of the log2(fold change) of the MANE (pink) and non-canonical isoforms (dark red) of genes that do not undergo significant alternative splicing following depletion of the indicated proteins.
**B)** Boxplots of the log2(fold change) of the MANE (teal) and non-canonical isoforms (dark green) of genes that undergo significant alternative splicing following depletion of the indicated proteins.
