## Supplemental Figure 5 for "Direct and indirect effects of spliceosome disruption compromise gene regulation by Nonsense-Mediated mRNA Decay"

### Figure S5

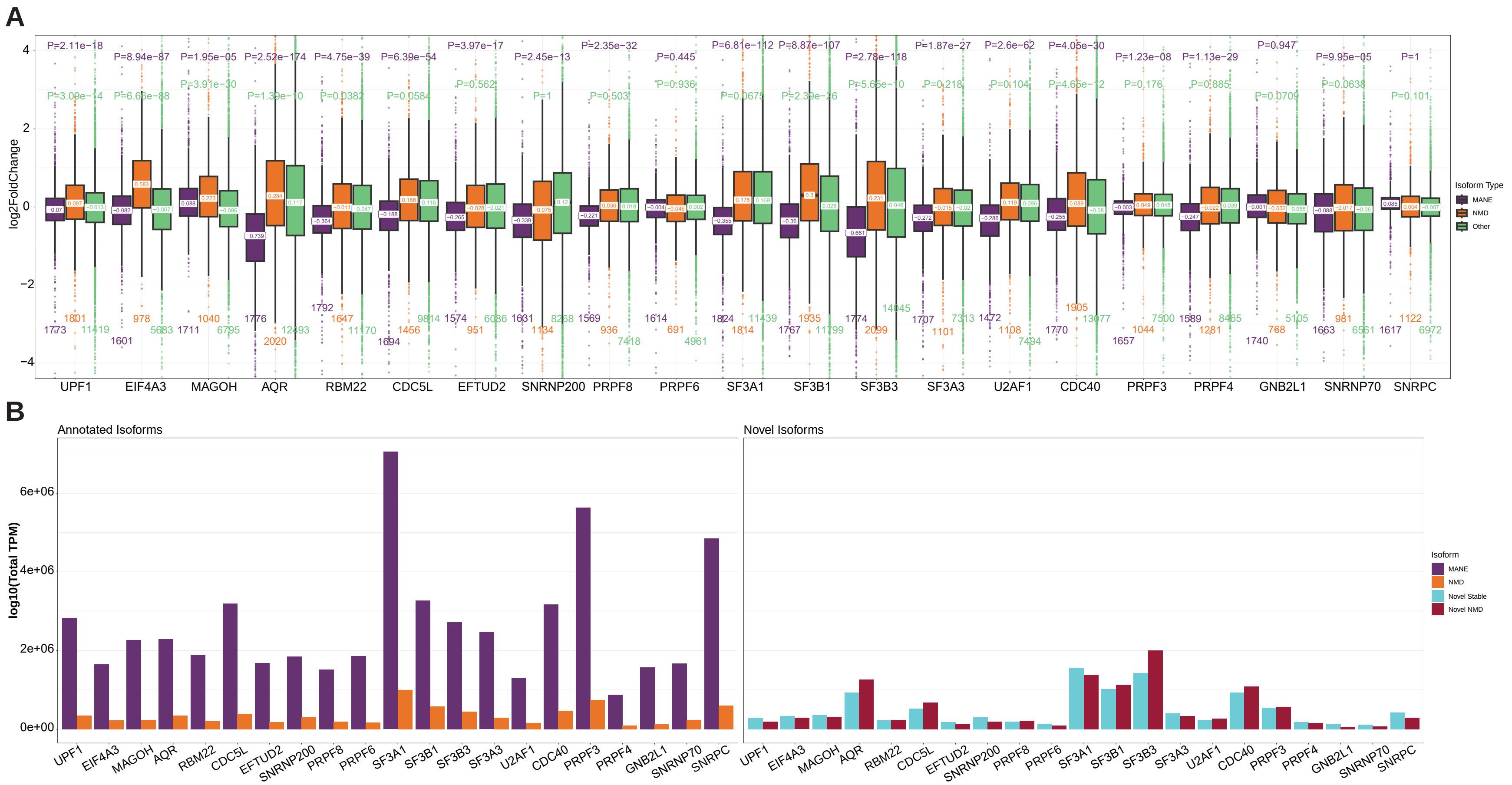


**Fig S5. Effect of spliceosome component depletion on relative levels of novel and annotated NMD targeted transcripts and of NMD factor mRNAs.**

**A)** Comparison of log2(fold change) of MANE (purple), NMD-targeted (orange), and stable non-canonical isoforms (green) following spliceosome component knockdown. The fold-changes were recalculated using kallisto and DESeq2 after including novel isoforms in the reference transcriptome. Median and number of observations in each group is noted as in Fig 2. P-value above the boxplots is the result of a Wilcoxon test comparing the MANE (purple) and stable non-canonical (green) isoforms to the PTC+ isoforms, with the alternative hypothesis being that the PTC+ isoforms will be more abundant.

**B)** Left: Cumulative length scaled TPM of MANE (purple) and NMD (orange) transcripts from genes that produce annotated NMD-targeted isoforms in the indicated samples. Right: Cumulative TPMs of stable (blue) and predicted NMD-targeted (red) novel isoforms produced from all genes in the indicated samples.
